# Sources of gene expression variation in a globally diverse human cohort

**DOI:** 10.1101/2023.11.04.565639

**Authors:** Dylan J. Taylor, Surya B. Chhetri, Michael G. Tassia, Arjun Biddanda, Alexis Battle, Rajiv C. McCoy

## Abstract

Genetic variation influencing gene expression and splicing is a key source of phenotypic diversity. Though invaluable, studies investigating these links in humans have been strongly biased toward participants of European ancestries, diminishing generalizability and hindering evolutionary research. To address these limitations, we developed MAGE, an open-access RNA-seq data set of lymphoblastoid cell lines from 731 individuals from the 1000 Genomes Project spread across 5 continental groups and 26 populations. Most variation in gene expression (92%) and splicing (95%) was distributed within versus between populations, mirroring variation in DNA sequence. We mapped associations between genetic variants and expression and splicing of nearby genes (*cis*-eQTLs and *cis*-sQTLs, respective), identifying >15,000 putatively causal eQTLs and >16,000 putatively causal sQTLs that are enriched for relevant epigenomic signatures. These include 1310 eQTLs and 1657 sQTLs that are largely private to previously underrepresented populations. Our data further indicate that the magnitude and direction of causal eQTL effects are highly consistent across populations and that apparent “population-specific” effects observed in previous studies were largely driven by low resolution or additional independent eQTLs of the same genes that were not detected. Together, our study expands understanding of gene expression diversity across human populations and provides an inclusive resource for studying the evolution and function of human genomes.

## Introduction

Genetic variation affecting gene expression and splicing accounts for a large proportion of phenotypic differences within and between species, including humans^1^. By correlating patterns of expression and splicing with variation at the level of DNA, foundational studies over the past decades have helped reveal the genetic architecture of these molecular traits, as well as their relationships to higher-order organismal phenotypes and diseases^2–6^.

Previous molecular association studies in humans have been strongly biased toward individuals of European ancestries, with limited representation from other global populations. This diminishes the generalizability of results, raises ethical concerns about the distribution of research benefits, and hinders understanding of the diversity and evolution of gene expression across human populations^7–9^. Further, research has demonstrated that the study of diverse cohorts captures a broader history of recombination events, improving the ability to identify the causal variant(s) driving an association signal against a background of correlated alleles in linkage disequilibrium (LD)^10^. More broadly, gene expression data from a large, globally diverse cohort is necessary to understand the distribution and evolution of human gene expression.

To address this goal, we developed MAGE: a resource for <u>M</u>ulti-ancestry <u>A</u>nalysis of <u>G</u>ene <u>E</u>xpression. MAGE comprises RNA-seq data from a large sample of lymphoblastoid cell lines from individuals from globally diverse human populations. Using these data, we quantified the distribution of gene expression and splicing diversity within and between populations, mapped genetic variation influencing gene expression and splicing at high resolution, and examined the evolutionary forces that shape such variation as well as the causes of apparent heterogeneity in its effects across populations. Together, our work offers a more complete view of the magnitude, distribution, and genetic sources of gene expression and splicing diversity within and between human populations.

## Results

### An open-access RNA-seq data set from globally diverse human samples

To achieve a broader view of human gene expression diversity, we performed RNA sequencing with poly(A) enrichment of lymphoblastoid cell lines from 731 individuals from the 1000 Genomes Project^11^ (1KGP), representing 26 globally-distributed populations (27-30 individuals per population) across five continental groups (**Fig. 1A**). These data offer a large, geographically diverse, open-access resource to facilitate studies of the distribution, genetic underpinnings, and evolution of variation in human transcriptomes and include data from several ancestry groups that were poorly represented in previous studies (**Fig. 1B, Fig. S2**). All 731 samples were sequenced in a single laboratory across 17 batches, and sample populations were stratified across these batches to avoid confounding between population and batch. In addition, 24 samples were sequenced in triplicate to facilitate quantification of technical variability both within and between batches, thus resulting in 779 total libraries.

**Figure 1.**
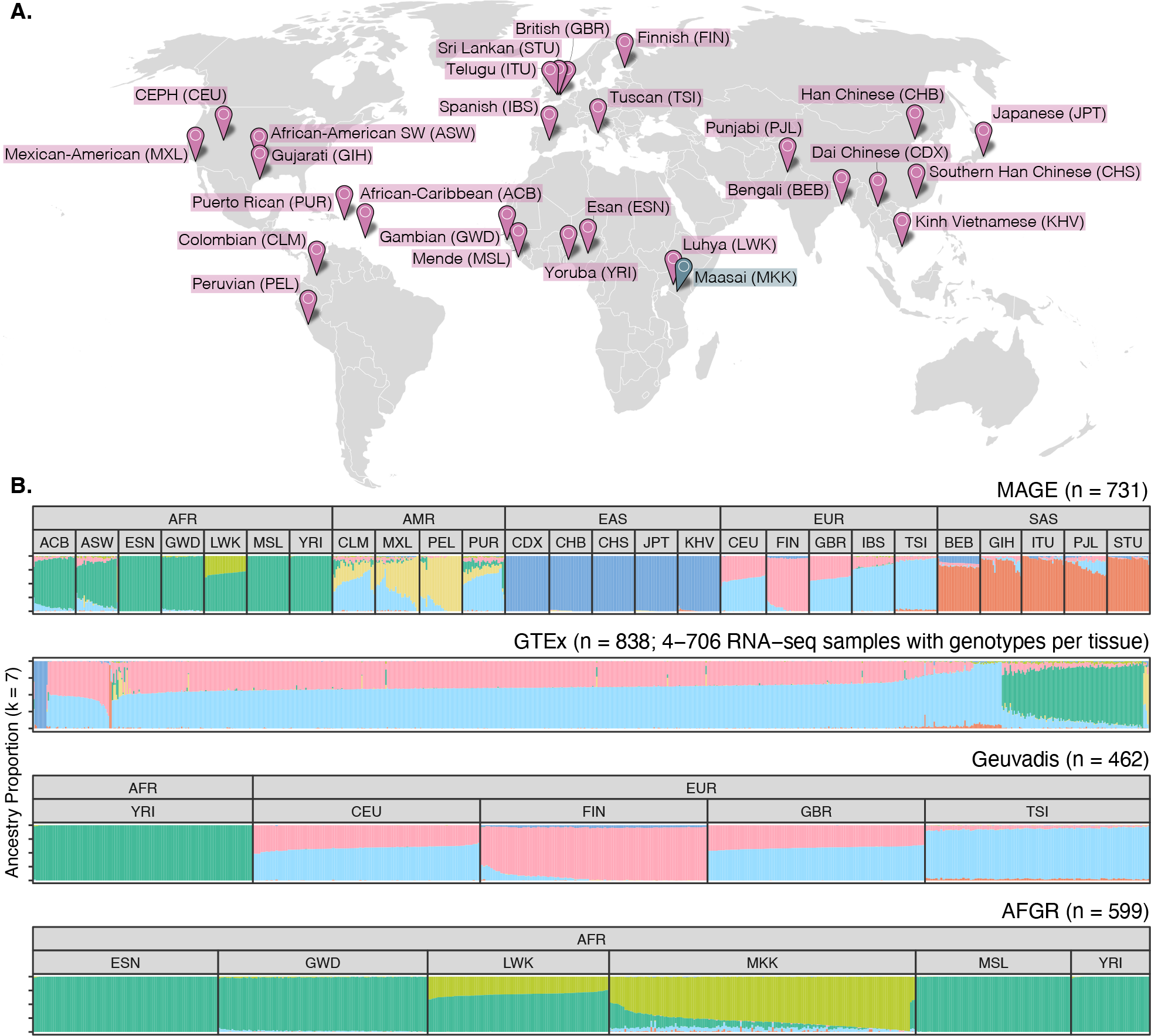
A globally diverse transcriptomics data set. **(A)** RNA-seq data was generated from lymphoblastoid cell lines from 731 individuals from the 1000 Genomes Project^11^, roughly evenly distributed across 26 populations and five continental groups. Populations included in MAGE are colored in pink, while the Maasai population is colored in blue as it is present in the AFGR^12^ data set (based on sequencing of HapMap^13^ cell lines), but not in the 1000 Genomes Project or MAGE. **(B)** ADMIXTURE^14^ results displaying proportions of individual genomes (columns) attributed to hypothetical ancestry components. For MAGE, Geuvadis^6^, and AFGR, samples are stratified according to population and continental group labels from the respective source projects, while GTEx^15^ does not include population labels.

We quantified gene expression level using gene annotations from GENCODE (v38; see Methods). After filtering out genes with low expression across samples, we were left with 20,154 expressed genes (19,539 autosomal genes, 615 genes on chrX) for downstream analysis of gene expression levels. In parallel, we used an annotation agnostic approach implemented by LeafCutter^16^ to quantify alternative splicing patterns, defined by ratios of intron excision events (see Methods). After filtering out low-count and low-complexity introns, we identified 32,867 splicing clusters (145,806 introns) mapping to 12,300 genes. Notably, for the 24 samples sequenced in triplicate, we observed greater variation in both gene expression and splicing between samples than between batches (**Fig. S3**), corroborating the robustness of our study design.

### The distribution of gene expression and splicing diversity

The vast majority of variation in DNA sequence is distributed within as opposed to between human populations by consequence of historical demographic events such as divergence, admixture, and population growth^17,18^. Previous studies have explored the extent to which this pattern holds for gene expression diversity, finding that population labels explain 3-25% of the total variation in gene expression^6,19^. However, these studies were limited by either sample size or diversity, motivating analysis within MAGE.

To this end, we fit a linear model relating expression level of each gene with continental group and population labels from the 1KGP. After regressing out sequencing batch and sex effects, continental group explains an average of 2.92% of variance in gene expression level across tested genes, whereas population label explains an average of 8.40% of variance, consistent with its more granular definition (**Fig. 2A**). While small, these proportions exceed null expectations assuming no population structure, supporting the interpretation that the geographic distribution of DNA sequence variation propagates to molecular phenotypes (permutation test: *p*_continental group_, *p*_population_ < 1 × 10^-3^). Interestingly, the proportion of variance explained is smaller, on average, than reported in a previous study by Martin et al.^19^. This may partially reflect their inclusion of samples from the San population of the Human Genome Diversity Project (HGDP) whose ancestors have been estimated to have separated (with subsequent gene flow) from other populations within their data set >100 kya, exceeding the greatest estimated divergence times between populations included in 1KGP^20^.

**Figure 2.**
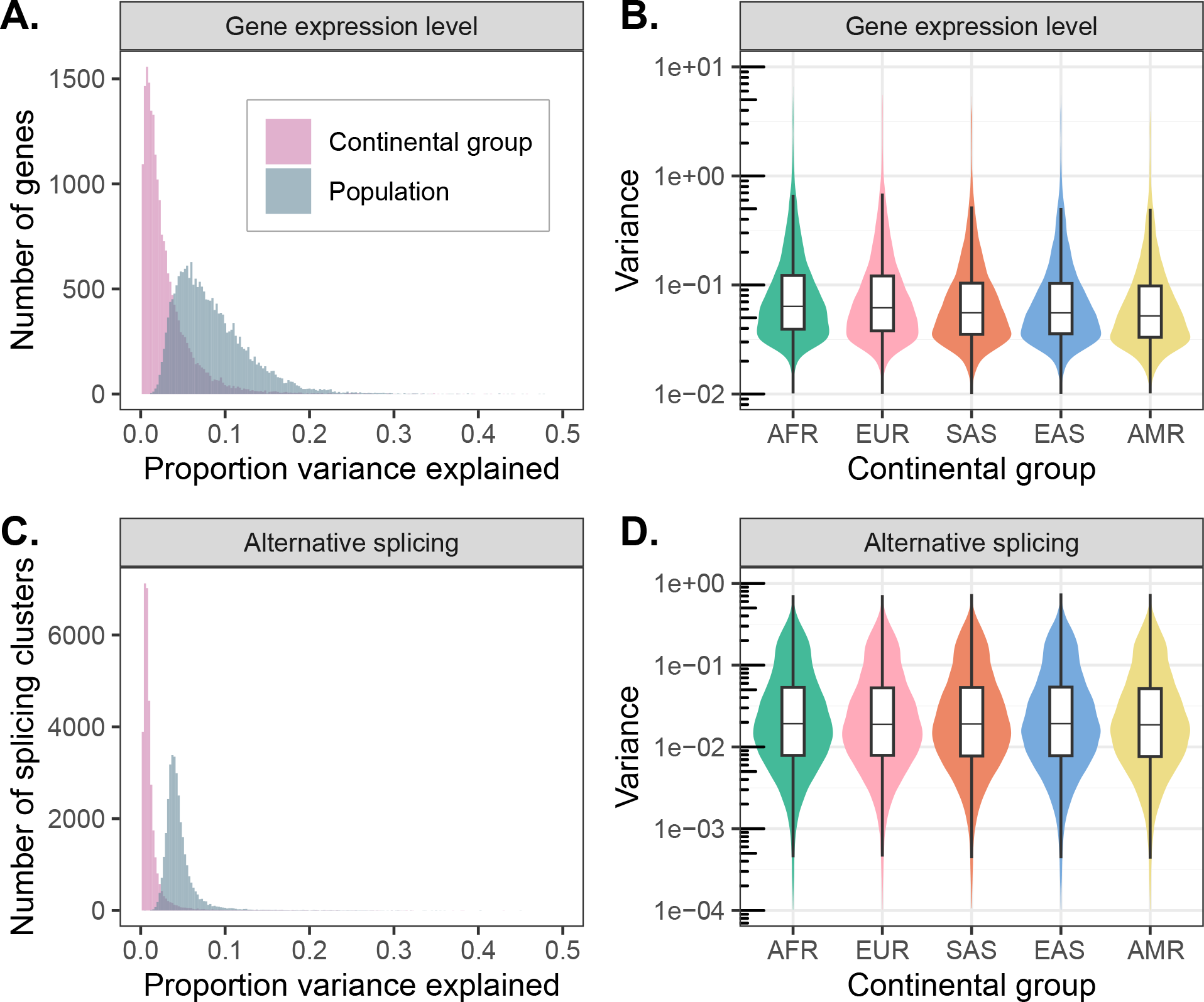
Patterns of transcriptomic diversity within and between populations. **(A)** Per gene estimates of the proportion of variance in gene expression level that is partitioned between continental groups (pink) and populations (blue), as opposed to within continental groups or populations. **(B)** Variance in gene expression level per gene differs across continental groups, consistent with underlying differences in levels of genetic variation. **(C)** Per splicing cluster estimates of the proportion of variance in alternative splicing (intron excision ratios) that is partitioned between continental groups (pink) and populations (blue), as opposed to within continental groups or populations. **(D)** Variance in alternative splicing (intron excision ratios) per splicing cluster differs across continental groups, consistent with underlying differences in levels of genetic variation.

We observed similar patterns of diversity across populations for alternative splicing. After regressing out technical variation, continental group and population explained an average of only 1.23% and 4.58%, respectively, of variance in splicing (**Fig. 2C**; permutation test: *p*_continental group_, *p*_population_ < 1 × 10^-3^). While the proportions of variance in gene expression and splicing explained by population label are not directly comparable due differences in their units of measurement (normalized expression levels per gene vs. intron excision ratios per splicing cluster), our observations are qualitatively consistent with previous reports that expression level varies more between populations than splicing^19^.

Interestingly, we also observed that within-population variance in expression (analysis of deviance: *χ*^2^ [4, *N* = 100,890] = 17,623, *p* < 1 × 10^-10^) and splicing (analysis of deviance: *χ*^2^ [4, *N* = 164,335] = 1550.6, *p* < 1 × 10^-10^) differs among continental groups, with higher average variances (across all tested genes) observed within the African continental group compared to the Admixed American continental group (**Fig. 2B and 2D, Fig. S5**). These results are consistent with the demonstrated decline in genetic diversity resulting from serial founder effects during human global migrations^21,22^. While significant, the magnitudes of these differences in variances are smaller than the magnitude of the decline in genetic diversity, likely reflecting the non-genetic environmental and stochastic contributions to gene expression and splicing variance that similarly affect all samples.

### Genetic effects on gene expression

#### Discovery of expression-linked genetic variation at high resolution

Beyond providing a global view of the patterns of expression and splicing across populations, MAGE offers a valuable resource for uncovering the genetic factors driving variation in these biological processes and the molecular mechanisms by which they act, including the discovery of functional variation that is largely private to historically understudied populations. By intersecting our expression and splicing quantifications with high-confidence genetic variant calls previously generated for the same set of samples^23^, we mapped expression and splicing quantitative trait loci within 1 megabase (Mb) of the transcription start site of each gene (termed *cis*-eQTLs and *cis*-sQTLs, respectively). We define eGenes and sGenes as genes with an eQTL or sQTL, respectively, and eVariants and sVariants as the individual genetic variants defining an eQTL or sQTL signal. Across 19,539 autosomal genes that passed expression level filtering thresholds (see Methods), we discovered 15,022 eGenes and 1,968,788 unique eVariants (3,538,147 significant eVariant– eGene pairs; 5% false discovery rate [FDR]). Additionally, across 11,912 autosomal genes that passed splicing filtering thresholds, we discovered 7,727 sGenes and 1,383,540 unique sVariants (2,416,177 significant sVariant–sGene pairs; 5% FDR).

The inclusion of genetically diverse samples in association studies captures a greater number of recombination events in the history of the sample, thereby breaking up linkage disequilibrium (LD) and improving mapping resolution^9,10^ (**Fig. S7**). With this goal in mind, we used SuSiE^24^ to perform fine-mapping for all eGenes and the introns of all sGenes to identify causal variants driving each QTL signal. For each gene/intron, SuSiE identifies one or more credible sets, representing independent causal e/sQTL signals and whereby each credible set contains as few variants as possible while maintaining a high probability of containing the causal variant. To obtain a gene-level summary of the sQTL fine-mapping results, we collapsed intron-level credible sets into gene-level credible sets by iteratively merging intron-level credible sets for each sGene (see Methods). We identified at least one credible set for 9,807 (65%) eGenes and 6,604 (85%) sGenes, which we define as fine-mapped e/sGenes. Consistent with previous results^6,15,25^, we observed widespread allelic heterogeneity across fine-mapped genes with 3,951 (40%) of fine-mapped eGenes and 3,490 (53%) of fine-mapped sGenes exhibiting more than one distinct credible set (**Fig. 3A, Fig. S9C**). We also achieved high resolution in identifying putative causal variants driving expression changes: of 15,664 eQTL credible sets, 3,992 (25%) contained just a single variant (median 5 variants per credible set; **Fig. 3B**). Similarly, for sQTLs, 3,569/16,451 (22%) credible sets contained just a single variant (median 7 variants per credible set; **Fig. S9D**). For downstream analyses, we selected a single representative “lead eQTL” from each eGene credible set, and a single “lead sQTL” from each sGene gene-level credible set (see Methods).

**Figure 3.**
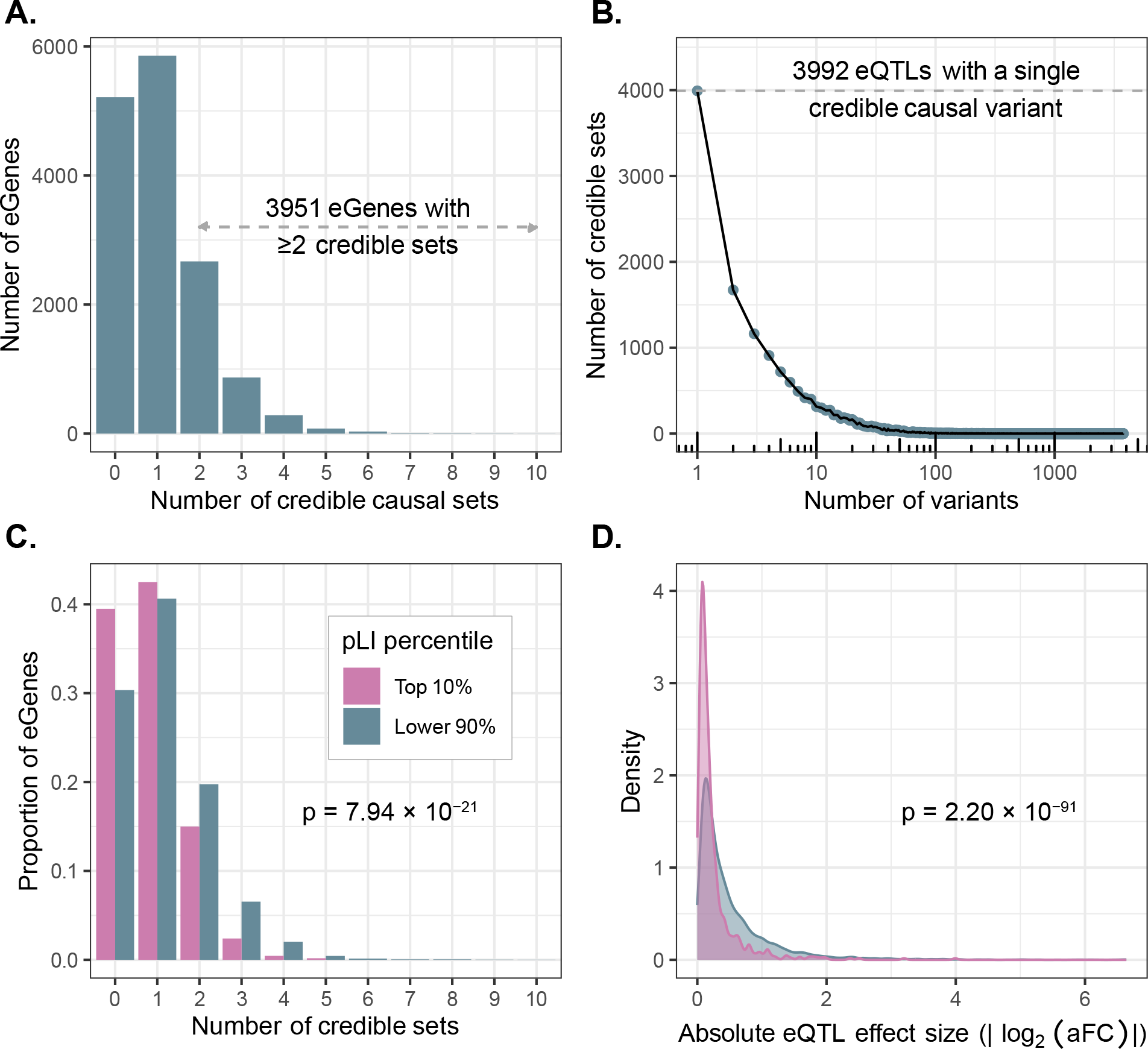
Mapping high-resolution eQTLs. **(A)** Number of credible sets per eGene, demonstrating evidence of widespread allelic heterogeneity whereby multiple causal variants independently modulate expression of the same genes. **(B)** Fine-mapping resolution, defined as the number of variants per credible set. **(C)** A signature of stabilizing selection on gene expression, whereby eGenes under strong evolutionary constraint (defined as the top pLI decile reflecting intolerance to loss of function mutations; pink) possess fewer credible sets, on average, than other genes (blue). **(D)** A signature of stabilizing selection on gene expression, whereby eQTLs of genes under strong evolutionary constraint (top pLI decile; pink) have smaller average effect sizes (aFC) than other genes (blue).

For each lead eQTL, we calculated its effect size using a recent implementation of the allelic fold change (aFC)^26^ statistic that quantifies eQTL effect sizes conditional on all other lead eQTLs for that gene (see Methods). Consistent with previous studies^15^, most eQTLs had small effects; only 2,031 (13%) lead eQTLs had greater than a two-fold effect on gene expression (median |log_2_(aFC)| = 0.30; **Fig. S8**). This is a slightly smaller proportion than previously reported by GTEx^15^, but we hypothesize that this is partially explained by small sample sizes in some GTEx tissues driving a stronger “winner’s curse” in those tissues, whereby effects are systematically overestimated^27^.

#### Evidence of selective constraint on eGenes

Previous studies of large population cohorts have identified sets of genes under strong mutational constraint, whereby negative selection has depleted loss-of-function point mutations and copy number variation relative to the rest of the genome^28^. One popular metric for quantifying mutational constraint on genes is the probability of intolerance to loss-of-function mutations (pLI)^28^. In our data, we observed that eGenes possess significantly lower mean pLI scores (mean = 0.261) than non-eGenes (mean = 0.304; Wilcoxon rank sum test: W = 11,596,590, *p* = 3.89 × 10^-7^). Additionally, highly constrained eGenes (top 10% of pLI) tend to possess fewer credible causal sets (mean = 0.82) than other eGenes (mean = 1.12; quasi-Poisson GLM: 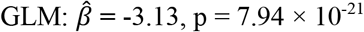 **Fig. 3C)**. Moreover, the average effect size of lead eQTLs within highly constrained genes (mean |log_2_(aFC)| = 0.26) is smaller than that of other genes (mean |log_2_(aFC)| = 0.53; Wilcoxon rank sum test: W = 4,128,766, *p* = 2.20 × 10^-91^; **Fig. 3D**). These observations, which hold for several other metrics of mutational constraint (**Fig. S10**), are consistent with previous analyses demonstrating weak, but measurable selection against expression-altering variation^29^.

#### Functional Enrichment of *cis*-eQTLs

Taking advantage of the high resolution of putative causal signals, we investigated the functional enrichment of fine-mapped *cis*-eQTLs in the regulatory DNA of multiple cell/tissue types. We quantified the enrichment of fine-mapped lead eQTLs in 15 predicted chromatin-state annotations across 127 reference epigenomes from the Roadmap Epigenomics chromHMM model^30^. We observed strong enrichment of lead eQTLs within several regulatory chromatin states (**Fig. 4A; S11A)** across cell/tissue types including within the blood and T-cell samples. Enrichment was most pronounced within promoter regions, specifically at active transcription start sites (TssA) and flanking regions (TssAFlnk), but modest enrichments are also apparent within enhancer regions (Enh, EnhG), especially for blood cell types. Conversely, quiescent, repressive, and heterochromatic regions were depleted of eQTLs. To parse cell-type-specific patterns, we further extended our analysis to primary DNAse Hypersensitivity Site (DHS) annotations, and we observed a strong enrichment of lead eQTLs in DHSs of blood and T-cell samples (**Fig. S11B**).

**Figure 4.**
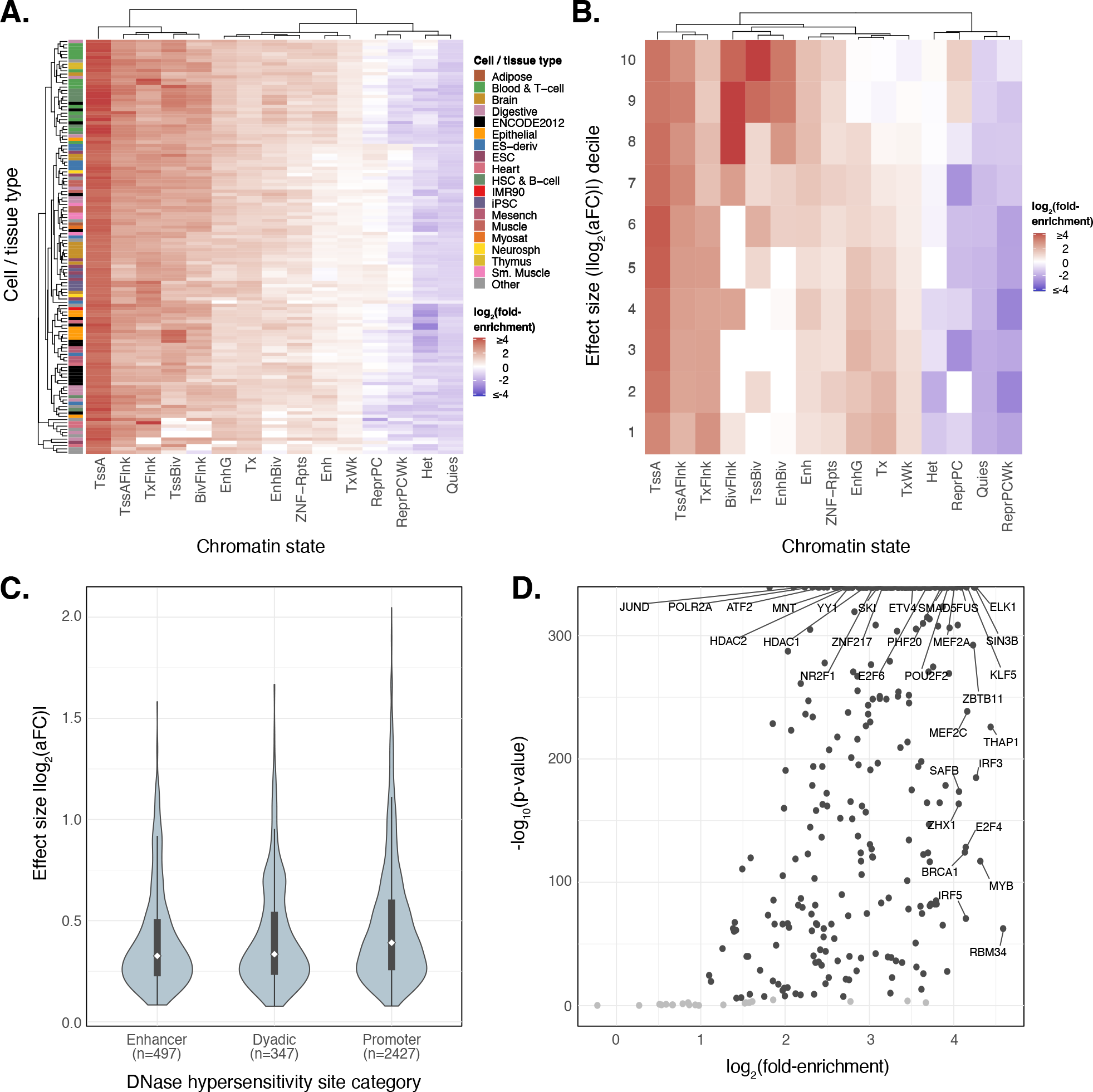
Fine-mapped *cis*-eQTLs are strongly enriched in regulatory regions across multiple cell/tissue types. **(A)** A heatmap representing hierarchical clustering of the enrichment of cis-eQTLs in predicted chromatin states using the Roadmap Epigenomics 15-state chromHMM model across 127 cell-tissue samples. **(B)** Enrichment analysis of the decile partitioned eQTL effect sizes measured as base-2 logarithm of the the absolute value of the estimated allelic fold change (|log_2_(aFC)|) across 15 different chromatin states predicted by chromHMM model specific to LCLs (Lymphoblastoid Cell Lines). **(C)** Distribution of absolute value of lead cis-eQTL effect sizes measured as log_2_(aFC) across putatively active chromatin states of LCLs linked to multi-tissue DNAse Hypersensitivity Sites. **(D)** Enrichment analysis of lead cis-eQTLs at TFBS (Transcription Factor Binding Sites) from ENCODE’s ChIP-seq binding profiles. Scatter data points with p-values < 0.001 (Bonferroni corrected) and log_2_(fold-enrichment) > 1 are colored in black underscoring those transcription factors where lead cis-eQTL enrichment is both statistically significant and of notable magnitude.

Focusing on data from LCLs, we next explored the relationship between epigenomic enrichments and eQTL effect sizes (|log_2_(aFC)|). While promoter-associated enrichment was consistent across eQTL effect size deciles, enrichment within poised regulatory regions such as Bivalent TSS (TSSBiv) and Bivalent Enhancers (EnhBiv) was most apparent for eQTLs of large effect (top two deciles), whereas quiescent, repressive, and heterochromatic regions again exhibited depletion (**Fig. 4B)**. Further exploring the effect sizes of functionally-annotated eQTLs, we observed that eQTLs located within chromatin states associated with transcriptional elongation regions (Tx, TxWk, and TxFlnk) predominantly exhibited lower effect sizes (**Fig. S12B**). These qualitative trends were replicated in other primary blood cell types (**Figs. S13-S16**).

To further contextualize the distribution of eQTLs effect sizes across different regulatory features, we analyzed the distribution of lead eQTL effect sizes (|log_2_(aFC)|) across a set of LCL promoter, enhancer, and dyadic (acting as both promoter and enhancer) regions from Roadmap Epigenomics that were annotated as active based on DHS signals across 53 cell/tissue types^30^. We observed larger median eQTL effect sizes at promoter regions relative to enhancers and dyadic regions (**Fig. 4C**)—a pattern that again replicates across other primary blood-related cell types (**Figs. S13-S16**). Using ChIP-seq data from ENCODE^31^, we also observed that lead-eQTLs are significantly enriched within 312 (92.30%; Bonferroni adjusted *p* < 0.05) transcription factor (TF) binding sites including canonical promoter-associated TFs such as POLR2A, TAF1, JUND, ATF2, and KLF5, as well as TFs such as HDACs, EP300, and YY1 which are more typically associated with enhancers (**Fig. 4D**). Our findings help illuminate the epigenomic features of gene regulation influenced by genetic variation, enhancing understanding of the functional consequences of *cis*-eQTLs in different cellular and chromatin contexts.

### Population-specific effects of genetic variation on gene expression

One fundamental question in association studies is the extent to which genetic associations replicate across human groups and the underlying factors driving heterogeneity between groups. Several previous studies have demonstrated that the predictive power of association study summary statistics (e.g., when used for the development of polygenic scores) declines when applied to groups whose ancestry does not match that of the discovery sample^8^. The underlying causes of such poor portability is a topic of active debate^32,33^, and several non-mutually exclusive explanations have been proposed: 1) differences in the allele frequency of causal variants between groups can lead to differential power to discover associations within those groups; 2) differences in patterns of LD between groups can lead to apparent nominal effect size heterogeneity of discovered variants between groups due to differences in LD either between a tag variant and the underlying causal SNP, or between multiple causal variants; 3) epistatic interaction between multiple causal SNPs, one or both of which exhibit differences in AF across groups can also lead to differences in the apparent nominal effect size between groups. Gene-by-environment interactions may also drive effect size heterogeneity, but we anticipate that such interactions are less relevant to our data given the common conditions used for deriving and culturing immortalized lymphoblastoid cell lines, as well as the block-randomized nature of the experimental design (see Methods).

To gain insight into the relative importances of these phenomena, we identified and characterized two broad classes of population-specific QTLs: 1) QTLs whose AF differs between continental groups (which we term frequency differentiated QTLs or fd-QTLs) and 2) QTLs that exhibit effect-size heterogeneity between continental groups (which we term heterogeneous effect QTLs or he-QTLs). We consider each class in the following sections.

#### Prevalence of population-stratified eQTLs

We hypothesized that the diversity of our sample would facilitate discovery of novel QTLs that are private to populations that were underrepresented in previous molecular association studies. To test this hypothesis, we evaluated the geographic frequency distribution of the 15,664 lead eQTLs for 9,807 genes with at least one credible set in MAGE. We defined the geographic frequency distribution as the variant frequency across continental regions (e.g., European [EUR]) defined in the sample. We observed that 8,837 (56.4%) lead eQTLs have an allele frequency greater than 5% across all regional groupings (i.e., “globally common”), consistent with the fact that statistical power for eQTL discovery scales with allele frequency and that most common variation is shared across human populations^34,35^ (**Fig. S17A and S17B**). However, we also identified 1,310 (8.3%) lead eQTLs across 1,210 unique genes that are unobserved in the European continental group, but present in one or more other continental groups, including 736 such variants that are common in Africa (**Fig. S17C**). An additional 115 (0.6%) lead eQTLs associated with 112 unique genes are unobserved in both European and African ancestry groups (**Fig. S17D**). Qualitatively similar patterns are also apparent for sQTLs (**Fig. S18**). The discovery of geographically restricted e- and s-QTLs that are beyond the major ancestries sampled by previous projects (e.g., GTEx and Geuvadis) further underscores the value of ancestrally diverse molecular QTL data sets.

To further contextualize our results, we compared our eQTL fine-mapping results to those from GTEx, which largely comprises individuals of European ancestries, as well as some African American subjects. To account for the multi-tissue nature of GTEx, we took the union of credible sets across tissues for a focal gene to compare with the credible sets for that same gene in MAGE (see Methods). Overall, we found that 8,069 MAGE credible sets (6,421 genes) replicate in GTEx, compared to 7,595 credible sets (5,545 genes) that do not replicate (**Fig. 5A**). We additionally identified 701 genes with at least one credible set in MAGE but no apparent credible set in GTEx. Notably, we observed that lead eQTLs in MAGE that do not replicate in GTEx tend to exhibit greater geographic differentiation with higher frequencies outside of Europe relative to variants that replicate between studies which tend to be common across all populations (**Fig. 5A**). Moreover, the 79,915 GTEx lead eQTLs that are not replicated in MAGE (7,913 lead eQTLs replicated) are enriched for tissue-specific effects (Mann-Whitney U Test: *p* < 10^-10^; **Fig. S19**). Importantly, the subset of MAGE eQTLs that do not replicate in GTEx exhibit similar qualitative patterns of functional enrichment as those that do replicate, thereby supporting the biological validity of newly discovered eQTLs in MAGE (**Fig. S20**). Together, these results highlight important aspects of experimental design across multiple axes of diversity, such as ancestry or tissue composition, that shape the statistical findings of molecular QTL studies.

**Figure 5.**
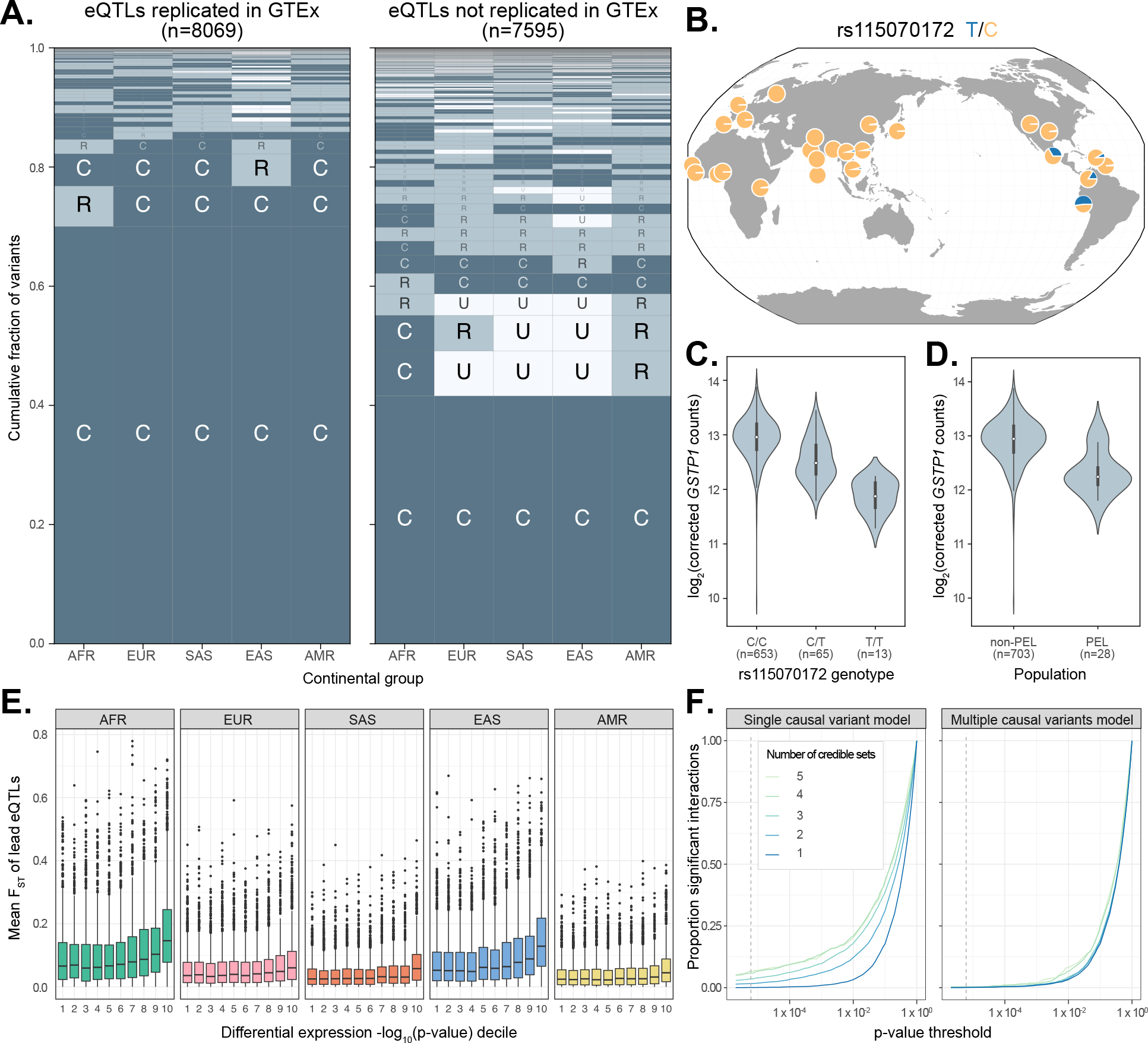
Population-specific genetic effects on gene expression. **(A)** Visualization of the joint distribution of allele frequencies of MAGE fine-mapped lead eQTLs across continental groups from the 1000 Genomes Project^11^, stratifying on replication status in GTEx^15^. **(B)** Allele frequency of a lead eQTL of *GSTP1* (rs115070172) across populations from the 1000 Genomes Project. The minor (T) allele achieves high frequency (AF = 0.63) in the Peruvian (PEL) population but is at moderate or low frequencies in other global populations. **(C)** Expression of *GSTP1*, stratified by genotype of rs115070172. **(D)** Expression of *GSTP1*, stratified by population label (PEL versus non-PEL). **(E)** Frequency differentiation (measured as mean F_ST_ between the focal continental group and all other groups) of lead eQTLs, stratifying by differential expression decile (contrasting expression in the focal continental group versus all other groups) of respective eGenes, where the 10th decile represents the strongest evidence of differential expression based on p-value. Across all continental groups, evidence of differential expression is positively associated with levels of frequency differentiation of lead eQTLs. **(F)** Number of significant genotype-by-continental group interactions at varying p-value thresholds (dashed line denotes Bonferroni threshold) for a model that considers a single causal variant at a time (left panel) versus a model that jointly considers multiple potential causal variants (right panel). Results are stratified by the number of credible sets (from one to five), demonstrating that failure to consider multiple causal variants may erroneously manifest as ancestry-specific heterogeneity of genetic effects on expression.

One example of an fd-eQTL that we identified was rs115070172, for which the T allele is common (AF > 0.05) only within the Admixed American (AMR) continental group in MAGE and is at high frequency (AF = 0.63) in the Peruvian (PEL) population (**Fig. 5B**). This variant is the lead eQTL for one of two credible causal sets of gene *GSTP1*, a tumor suppressor gene whose expression has been implicated in breast cancer^36–38^. The T allele of the rs115070172 variant is significantly associated with lower expression of *GSTP1* (**Fig. 5C**). Intersection with epigenomic data indicate that this fine-mapped lead eQTL lies within a putative enhancer region. We note that there is also a second SNP, rs4930437, in the same credible set that is completely linked with the chosen lead eQTL and lies within a putative promoter region and is proximal (471 bp) to an H3K27ac-enriched site, a hallmark of strong enhancer activity (**Fig. S21**). Interestingly, the expression of *GSTP1* is significantly lower in individuals from PEL compared to other global populations, and we hypothesize that this eQTL signal may explain this trend.

To more broadly test the role of fd-eQTLs in driving differential gene expression between continental groups, we quantified Weir & Cockerham’s F^ST39^ for each lead eQTL and intersected these values with the differential expression (DE) results for each eQTL’s respective eGene (**Fig 5E**). Among continental groups, differentially expressed eGenes (FDR-adjusted *p* ≤ 0.05) possessed higher F_ST_ values than non-DE genes (Mann-Whitney U Test: Z = 0.022 ± 0.001, 95% c.i.; *p* < 0.05). Within continental groups, the magnitude of differential expression positively correlated with mean F_ST_, where the greatest difference in F_ST_ was measured within AFR (mean F_ST_ = 9.92 × 10^-2^ in 1^st^ DE decile; mean F_ST_ = 1.82 × 10^-1^ in 10^th^ DE decile) and the smallest difference within AMR (mean F_ST_ = 3.76 × 10^-2^ in 1^st^ DE decile; mean F_ST_ = 6.37 × 10^-1^ in 10^th^ DE decile). This result suggests that gene expression differences across populations can be attributed to frequency differentiation of causal eQTLs.

#### eQTL effect sizes are consistent across continental groups

Given the debate regarding the prevalence of GWAS/eQTL hits exhibiting effect-size heterogeneity between ancestries/populations^9,32,33,40,41^, we sought to test for he-eQTLs in MAGE. Because the genotypes are derived from high-coverage whole genome sequencing in 1KGP, MAGE should be robust to effect size heterogeneity resulting from population-specific LD patterns with an untyped casual variant (as commonly affects microarray data), barring large structural variation that may escape detection with short-read sequencing. This allows us to investigate other potential sources of effect size heterogeneity. For each fine-mapped lead eQTL, we compared the standard (i.e., nominal pass) association model to a model that includes an additional genotype-by-continental group interaction term, testing whether the more complex interaction model explains the data significantly better than the reduced model. Across 12,338 lead eQTLs that passed filtering (MAF ≥ 0.05 in at least two continental groups), 204 (1.65%) had a significant interaction with continental group after Bonferroni correction (**Fig. 5F**).

Interestingly, eGenes with more credible sets were more likely to exhibit significant genotype-by-continental group interactions, suggesting that the additive effects of multiple causal variants may drive apparent interaction effects. To test this hypothesis, we compared a second set of models that, for each lead eQTL, include all other lead eQTLs for that gene. Supporting our hypothesis, 198 (97%) of lead eQTLs that had a significant interaction effect lost significance after controlling for the additive effects of the other causal signals for that gene (**Fig. 5F**). This suggests that effect size heterogeneity of eQTLs between populations is rare, and apparent heterogeneity may instead reflect the failure to control for the additive effects of multiple independent causal signals. This conclusion is consistent with previous studies using orthogonal approaches for evaluating effect size heterogeneity based on analysis of admixed individuals^33,40^. For the small number of lead eQTLs that do have a significant interaction effect after controlling for multiple causal signals (9 eQTLs; 0.07% of all eQTLs that passed filtering), this effect could be driven by non-additive epistatic interactions between variants, additional untested causal variants that did not meet nominal MAF thresholds, or population-specific LD patterns with a untyped causal variants. This result suggests that *trans*-genetic effects (driven by global ancestry patterns), if properly controlled for, generally do not have a strong impact on the effects of causal variants in *cis*. Further, this is encouraging for predictive applications such as polygenic risk scores (PRS) and transcriptome-wide association studies (TWAS), as it suggests that properly constructed models that 1) focus on causal signals and, 2) do not make assumptions about the number of such signals may exhibit better portability between groups.

## Discussion

Combined with existing whole-genome sequencing data from the same set of samples^23^, MAGE offers a large open-access data set for studying the diversity and evolution of human gene expression and splicing. Our study also offers powerful insight into the genetic sources of variation in these key molecular phenotypes which may in turn modulate organismal phenotypes (though see reference^42^ for statistical insight into the limited overlap between molecular QTLs and GWAS hits). By evenly spanning samples from all 26 populations of 1KGP^11^, MAGE includes several ancestry groups that were poorly represented in previous molecular association studies, thereby addressing a long-standing bias within the field of human genetics^7^.

The scale and diversity of the data set enabled the discovery of numerous potentially novel genetic associations, while also offering high resolution for identifying putative causal variants and elucidating their mechanisms of action. Our study also demonstrates that conditional on the correct identification and presence of causal variants, the effects of such variants tend to be additive and highly consistent across populations—a point of recent debate within the field^9,32,33,40,41^. This observation in turn suggests that ancestry-dependent epistatic effects tend to be weak and/or rare in human genomes, in contrast to some observations from other model systems^43^. Such consistency of genetic effects further motivates the use of diverse samples for association studies, as a common causal variant identified in one population may inform the effect of that variant in a population where the same variant is rare and association testing would be underpowered.

By design, our study focuses on a single cell type of lymphoblastoid cell lines, which offer a useful model for studying gene expression given their low somatic mutation rates and robust gene expression patterns encompassing key metabolic pathways^44^. While this allows us to mitigate the effects of environmental variation and compare our results to related studies performed in the same cell lines^6^, future studies may seek to understand ancestry differences in expression across developmental, cellular, and other environmental contexts, including with respect to dynamic QTLs whose effects vary based on those contexts^45^. Future studies of diverse cohorts may also leverage new technologies (such as long-read genome, cDNA, or direct RNA sequencing^46–48^) and/or novel computational approaches such as those based on pangenomes^49^ to achieve higher resolution for isoform detection as well as improved analysis of genes occurring within highly repetitive or structurally complex regions. Finally, while geographically diverse, the sampling of 1KGP is not without biases—for example, narrowly sampling the vast diversity within Africa and excluding indigenous populations from Oceania and the Americas, as well as countless other populations. Addressing these biases will require deeper community engagement and respect for the rights, interests, and expectations of research participants from diverse human groups^50^.

Our work offers a more complete picture of the links between genetic variation and genome function across diverse populations, as well as the evolutionary forces that have shaped this variation within our species. Complemented by existing high-coverage whole genome sequencing data, we anticipate that this data set will serve as a valuable resource to facilitate future research into the complex genetic basis of variation in human genome function.

## Data availability

Newly generated RNA sequencing data for the 731 individuals (779 total libraries) is available on the Sequence Read Archive (Accession: PRJNA851328). Processed gene expression matrices and QTL mapping results are available through Dropbox as described on GitHub at https://github.com/mccoy-lab/MAGE.

## Code availability

Code used for the analyses presented in this paper is available at https://github.com/mccoy-lab/MAGE.

## Acknowledgments

Thank you to Stephen Montgomery and the Montgomery lab, Genevieve Wojcik, Jeff Leek, and members of the McCoy Lab, Department of Biology, and Center for Computational Biology at Johns Hopkins for helpful discussion and feedback. Thank you to Calvin Runnels for initial exploration of signatures of selection. We also thank the staff of Advanced Research Computing at Hopkins for support, as well as Matthew Mitchell and other staff at the Coriell Institute, Patrick Boyle, Brittany Kerr, Ashley James, Aline Bronzato-Badial, McKenzie Carter, and other staff at Genewiz for assistance generating RNA sequencing data. RCM, MGT, and A. Biddanda are supported by NIH/NIGMS Award R35GM133747. DJT is supported by NIH/NHGRI Award F31HG012900. A. Battle is supported by NIH/NIGMS Award R35GM139580. The content is solely the responsibility of the authors and does not necessarily represent the official views of the National Institutes of Health.

## References

1. Li, Y. I. et al. RNA splicing is a primary link between genetic variation and disease. Science 352, 600–604 (2016).

2. Brem, R. B., Yvert, G., Clinton, R. & Kruglyak, L. Genetic dissection of transcriptional regulation in budding yeast. Science 296, 752–755 (2002).

3. Morley, M. et al. Genetic analysis of genome-wide variation in human gene expression. Nature 430, 743–747 (2004).

4. GTEx Consortium. Genetic effects on gene expression across human tissues. Nature 550, 204–213 (2017).

5. Mogil, L. S. et al. Genetic architecture of gene expression traits across diverse populations. PLoS Genet. 14, e1007586 (2018).

6. Lappalainen, T. et al. Transcriptome and genome sequencing uncovers functional variation in humans. Nature 501, 506–511 (2013).

7. Popejoy, A. B. & Fullerton, S. M. Genomics is failing on diversity. Nature 538, 161–164 (2016).

8. Martin, A. R. et al. Human Demographic History Impacts Genetic Risk Prediction across Diverse Populations. Am. J. Hum. Genet. 107, 788–789 (2020).

9. Wojcik, G. L. et al. Genetic analyses of diverse populations improves discovery for complex traits. Nature 570, 514–518 (2019).

10. Kita, R., Venkataram, S., Zhou, Y. & Fraser, H. B. High-resolution mapping of cis-regulatory variation in budding yeast. Proc Natl Acad Sci U S A . 114, E10736–E10744 (2017).

11. 1000 Genomes Project Consortium et al. A global reference for human genetic variation. Nature 526, 68–74 (2015).

12. DeGorter, M. K. et al. Transcriptomics and chromatin accessibility in multiple African population samples. bioRxiv.

13. International HapMap Consortium. The International HapMap Project. Nature 426, 789–796 (2003).

14. Alexander, D. H., Novembre, J. & Lange, K. Fast model-based estimation of ancestry in unrelated individuals. Genome Res. 19, 1655–1664 (2009).

15. GTEx Consortium. The GTEx Consortium atlas of genetic regulatory effects across human tissues. Science 369, 1318–1330 (2020).

16. Li, Y. I. et al. Annotation-free quantification of RNA splicing using LeafCutter. Nat. Genet. 50, 151–158 (2018).

17. Lewontin, R. C. The apportionment of human diversity. in Evolutionary Biology 381–398 (Springer US, 1972).

18. Jorde, L. B. et al. The distribution of human genetic diversity: a comparison of mitochondrial, autosomal, and Y-chromosome data. Am. J. Hum. Genet. 66, 979–988 (2000).

19. Martin, A. R. et al. Transcriptome sequencing from diverse human populations reveals differentiated regulatory architecture. PLoS Genet. 10, e1004549 (2014).

20. Bergström, A. et al. Insights into human genetic variation and population history from 929 diverse genomes. Science 367, (2020).

21. Ramachandran, S. et al. Support from the relationship of genetic and geographic distance in human populations for a serial founder effect originating in Africa. Proc. Natl. Acad. Sci. U. S. A. 102, 15942–15947 (2005).

22. Prugnolle, F., Manica, A. & Balloux, F. Geography predicts neutral genetic diversity of human populations. Curr. Biol. 15, R159–60 (2005).

23. Byrska-Bishop, M. et al. High-coverage whole-genome sequencing of the expanded 1000 Genomes Project cohort including 602 trios. Cell 185, 3426–3440.e19 (2022).

24. Zou, Y., Carbonetto, P., Wang, G. & Stephens, M. Fine-mapping from summary data with the ‘Sum of Single Effects’ model. PLoS Genet. 18, e1010299 (2022).

25. Jansen, R. et al. Conditional eQTL analysis reveals allelic heterogeneity of gene expression. Hum. Mol. Genet. 26, 1444–1451 (2017).

26. Mohammadi, P., Castel, S. E., Brown, A. A. & Lappalainen, T. Quantifying the regulatory effect size of cis-acting genetic variation using allelic fold change. Genome Res. 27, 1872–1884 (2017).

27. Huang, Q. Q., Ritchie, S. C., Brozynska, M. & Inouye, M. Power, false discovery rate and Winner’s Curse in eQTL studies. Nucleic Acids Res. 46, e133 (2018).

28. Lek, M. et al. Analysis of protein-coding genetic variation in 60,706 humans. Nature 536, 285–291 (2016).

29. Glassberg, E. C., Gao, Z., Harpak, A., Lan, X. & Pritchard, J. K. Evidence for Weak Selective Constraint on Human Gene Expression. Genetics 211, 757–772 (2019).

30. Roadmap Epigenomics Consortium et al. Integrative analysis of 111 reference human epigenomes. Nature 518, 317–330 (2015).

31. ENCODE Project Consortium. An integrated encyclopedia of DNA elements in the human genome. Nature 489, 57–74 (2012).

32. Patel, R. A. et al. Genetic interactions drive heterogeneity in causal variant effect sizes for gene expression and complex traits. Am. J. Hum. Genet. 109, 1286–1297 (2022).

33. Hou, K. et al. Causal effects on complex traits are similar for common variants across segments of different continental ancestries within admixed individuals. Nat. Genet. 55, 549–558 (2023).

34. Visscher, P. M. et al. 10 Years of GWAS Discovery: Biology, Function, and Translation. Am. J. Hum. Genet. 101, 5–22 (2017).

35. Gutenkunst, R. N., Hernandez, R. D., Williamson, S. H. & Bustamante, C. D. Inferring the Joint Demographic History of Multiple Populations from Multidimensional SNP Frequency Data. PLoS Genet. 5, (2009).

36. Fang, C. et al. Aberrant GSTP1 promoter methylation is associated with increased risk and advanced stage of breast cancer: a meta-analysis of 19 case-control studies. BMC Cancer 15, 1–8 (2015).

37. Louie, S. M. et al. GSTP1 Is a Driver of Triple-Negative Breast Cancer Cell Metabolism and Pathogenicity. Cell Chemical Biology 23, 567–578 (2016).

38. Arai, T. et al. Association of GSTP1 CpG Islands Hypermethylation with Poor Prognosis in Human Breast Cancers. Breast Cancer Res. Treat. 100, 169–176 (2006).

39. Weir, B. S. & Cockerham, C. C. ESTIMATING F-STATISTICS FOR THE ANALYSIS OF POPULATION STRUCTURE. Evolution 38, 1358–1370 (1984).

40. Saitou, M., Dahl, A., Wang, Q. & Liu, X. Allele frequency differences of causal variants have a major impact on low cross-ancestry portability of PRS. medRxiv 2022.10.21.22281371 (2022) doi:10.1101/2022.10.21.22281371.

41. Kachuri, L. et al. Gene expression in African Americans, Puerto Ricans and Mexican Americans reveals ancestry-specific patterns of genetic architecture. Nat. Genet. 55, 952–963 (2023).

42. Mostafavi, H., Spence, J. P., Naqvi, S. & Pritchard, J. K. Systematic differences in discovery of genetic effects on gene expression and complex traits. Nat. Genet. (2023) doi:10.1038/s41588-023-01529-1.

43. Rau, C. D. et al. Modeling epistasis in mice and yeast using the proportion of two or more distinct genetic backgrounds: Evidence for ‘polygenic epistasis’. PLoS Genet. 16, e1009165 (2020).

44. Cheung, V. G. et al. Natural variation in human gene expression assessed in lymphoblastoid cells. Nat. Genet. 33, 422–425 (2003).

45. Strober, B. J. et al. Dynamic genetic regulation of gene expression during cellular differentiation. Science 364, 1287–1290 (2019).

46. Workman, R. E. et al. Nanopore native RNA sequencing of a human poly(A) transcriptome. Nat. Methods 16, 1297–1305 (2019).

47. Glinos, D. A. et al. Transcriptome variation in human tissues revealed by long-read sequencing. Nature 608, 353–359 (2022).

48. Reese, F. et al. The ENCODE4 long-read RNA-seq collection reveals distinct classes of transcript structure diversity. bioRxiv (2023) doi:10.1101/2023.05.15.540865.

49. Sibbesen, J. A. et al. Haplotype-aware pantranscriptome analyses using spliced pangenome graphs. Nat. Methods 20, 239–247 (2023).

50. Claw, K. G. et al. A framework for enhancing ethical genomic research with Indigenous communities. Nat. Commun. 9, 2957 (2018).

