## Supplementary Materials for "Sources of gene expression variation in a globally diverse human cohort"

Taylor *et al.*, 2023


**Supplemental Methods:** **3**

Study design 3

Ancestry analysis 3

RNA sequencing data production 5

Cell line processing and shipping 5

RNA extraction and sequencing 5

Preliminary gene expression level quantification 5

Gene-level counts 5

Lowly-expressed gene filter 5

Preliminary alternative splicing quantification 6

Quantification of intron excision ratios 6

Filtering lowly-expressed and low-complexity clusters 6

Quantifying the contribution of batch effects to expression variation 7

Differential gene expression between populations 9

Data preparation 9

Factor contrasts 9

Expression level variation within and between populations 9

Normalized expression matrix 9

Estimation of biological variation 10

Splicing variation within and between populations 11

*cis*-eQTL mapping 12

Expression normalization 12

Calculation of genotype PCs 12

Calculation of PEER covariates 13

Discovery of nominal *cis*-eQTLs with FastQTL 14

Fine-mapping eGene credible sets with SuSiE 14

Comparison of fine-mapping resolution in subsets of MAGE 15

Calculation of Allelic Fold Change (aFC) 16

*cis*-sQTL mapping 17
Splicing normalization 17
Calculation of PEER covariates 17
Discovery of nominal *cis*-sQTLs with FastQTL 17
Fine-mapping sGene credible sets with SuSiE 17
Analysis of negative selection 19
Functional annotation and enrichment of fine-mapped *cis*-QTLs 20
Lead e- and sQTL AF differentiation between populations 27
Replication of credible sets in GTEx 28
Defining replicating vs. non-replicating eQTLs 28
Functional annotation and enrichment of non-replicating eQTLs 29
Relationship between fixation index and differential gene expression 32
*cis*-eQTL effect size heterogeneity between populations 32
**References:** 34

### Supplemental Methods:

#### 1 Study design

RNA sequencing was performed entirely by GENEWIZ, from Azenta Life Sciences in South Plainfield, New Jersey. We performed RNA sequencing of 779 cell lines. These cell lines represent 731 unique samples, 24 of which were sequenced in triplicate. Sequencing was performed in batches of 15-48 cell lines each (twelve batches of 48 cell lines, four batches of 47 cell lines, and one batch of 15 cell lines). For samples with replicates, replicates were divided between batches such that one replicate of the three was sequenced in one batch, and the other two replicates were sequenced in a separate batch to allow for analysis of inter- and intra-batch variation for each of these samples.

#### 2 Ancestry analysis

Ancestry composition of our study sample was assessed and compared to related studies using ADMIXTURE[14], which uses a likelihood model to estimate allele frequencies in *k* postulated ancestral populations, as well as ancestry proportions for each individual that trace to each of those *k* populations. Genotype data for 1000 Genomes Project[11] (1KGP) samples were obtained from published data based on high coverage (~30×) sequencing by the New York Genome Center (NYGC)[23], subsetting to samples used in our study, by the Geuvadis consortium[6], and or the African Functional Genomics Resource[12] (AFGR). Data were downsampled to SNPs in approximate linkage equilibrium (using the `--indep-pairwise 200 20 0.2` flag in PLINK[51]) and restricted to common variants with MAF > 0.05 within the sample. This set of variants was then extracted from genotype data from v8 of the GTEx Project[15] as well as genotype data for samples from the Maasai (MKK) population from AFGR (which are not part of 1KGP), requiring that the SNPs be polymorphic and biallelic (with the same two alleles) in all data sets. Genotype data from this subset of variants was then merged across the relevant data sets and used as input to ADMIXTURE with default stopping criteria. For the purpose of visualization, *k* was set to 7, which exhibited the minimum 5-fold cross-validation error for the tested range of *k* = [2 .. 10] (**Fig. S1**). Principal components analysis was performed on the same merged data set using PLINK[51] (**Fig. S2**).

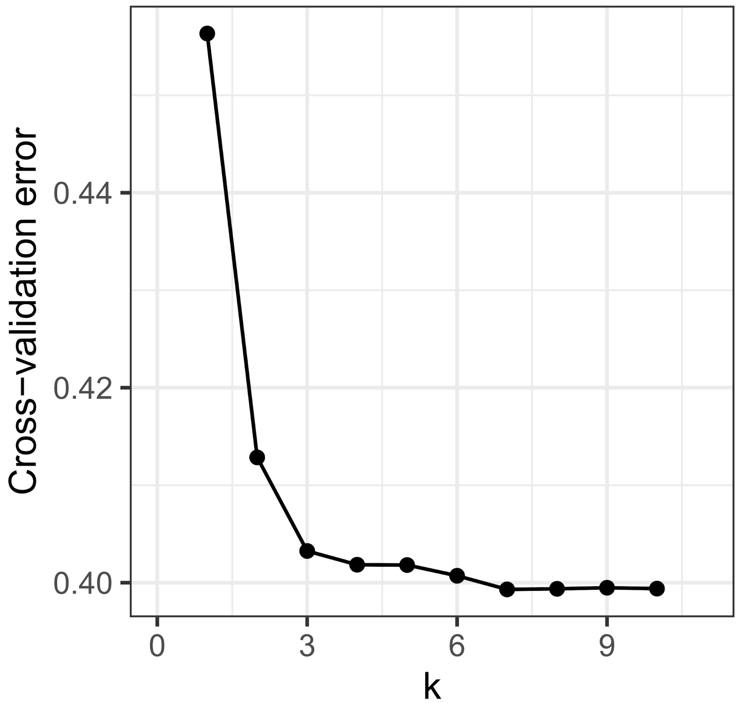


**Figure S1. ADMIXTURE cross validation error.** Five-fold cross-validation error with varying numbers of specified ancestry components (k) in ADMIXTURE. We selected k=7 for use in Figure 1 as this value minimizes the cross-validation error.

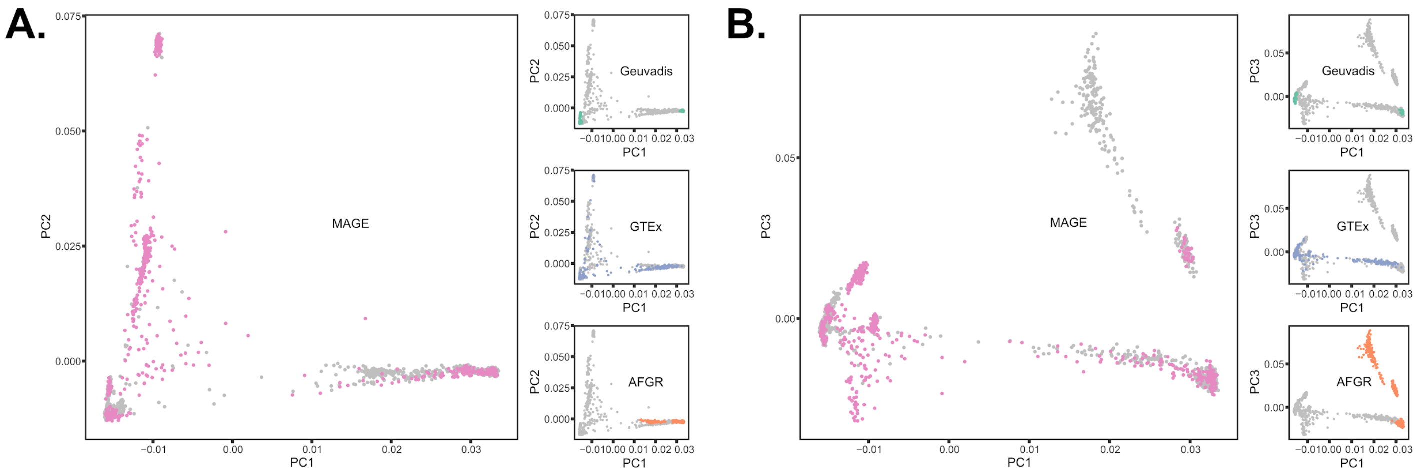


**Figure S2. Principal components analysis of genotype data corresponding to various human RNA-sequencing genomic data sets. (A)** Genotype principal components 1 and 2 with samples from all studies depicted with gray points and samples from the specified study (i.e., MAGE [pink], Geuvadis [green], GTEx [blue], and AFGR [orange]) depicted with colored points in each respective panel. **(B)** Same as panel A, but for principal components 1 and 3.

### 3 RNA sequencing data production

#### 3.1 Cell line processing and shipping

EBV transformed lymphoblastoid cells lines (LCLs) were purchased from the Coriell Institute for Medical Research (NIGMS and NHGRI Repositories) in Camden, New Jersey. Frozen cell pellets (≥ 5 million cells per cell line) were recovered by Coriell and cultured for 4 days (see Coriell LCL culture FAQ for information about growth media: www.coriell.org/0/sections/support/global/Lymphoblastoid.aspx). After growth, cells were transferred to a growth-limiting shipping media and were shipped directly to GENEWIZ (same-day delivery) for RNA isolation, library prep, and sequencing.

#### 3.2 RNA extraction and sequencing

At CORIELL, cells were spun down, then total RNA was extracted from cell pellets using Qiagen RNeasy Plus Universal mini kit following manufacturer's instructions. RNA was quantified using Qubit 2.0 Fluorometer and RNA integrity was checked using the Agilent TapeStation. Sequencing libraries were prepared using the unstranded NEBNext Ultra II RNA Library Prep Kit for Illumina using manufacturer's instructions with the polyA enrichment workflow. Sequencing libraries were validated on the Agilent TapeStation and quantified by using Qubit 2.0 Fluorometer as well as by quantitative PCR. Sequencing was done on the Illumina NovaSeq 6000 instrument with 150bp paired-end sequencing, with a desired minimum depth of 25M reads per sample. Sequencing libraries were multiplexed in batches of 15-48 samples (the same batches they were shipped in) and loaded onto the flow cell according to manufacturer's instructions.

### 4 Preliminary gene expression level quantification

#### 4.1 Gene-level counts

To quantify gene expression level in our data set, we use the GENCODE v38 transcript annotations[52] and Salmon (version 1.5.2)[53] for expression quantification. Salmon is a kmer-based method that uses raw RNA-seq data to estimate the number of reads aligning to a defined set of transcripts and their relative abundance. We first generated a Salmon index using `salmon index` with the GENCODE v38 transcript FASTA file as input and with the `--gencode` flag. For each of the 779 cell lines, we quantify transcript-level expression using `salmon quant` with the raw RNA-seq reads as input. We set `--libType=IU` because our sequencing pipeline is expected to produce read-pairs that are inwardly-oriented and unstranded. All other arguments use their default values. This produces, for each library, transcript-level estimates of read counts and TPM. Finally, these transcript-level estimates were summed to gene-level estimates using tximport (version 1.18.0)[54] in R. These gene-level quantifications are used as a starting point in down-stream analyses. Unless otherwise stated, for the 24 samples that were sequenced in triplicate, downstream analyses are limited to the replicate with the most reads for each of these samples.

#### 4.2 Lowly-expressed gene filter

For most analyses of gene expression level differences, it is useful to filter out genes with low expression across samples. Expression quantifications for lowly-expressed genes may be indistinguishable from sequencing noise and can introduce false-positive results across analyses. As such, we limit most analyses to genes with ≥6 counts and ≥0.1 TPM in at least 20% (147/731) of samples. After filtering, we were left with 20,154 expressed genes (19,539 autosomal genes, 615 genes on chrX) used for analyses of gene expression level.

### 5 Preliminary alternative splicing quantification

#### 5.1 Quantification of intron excision ratios

To quantify alternative splicing in our data set, we followed the splicing quantification pipeline developed by the GTEx consortium and described in their paper[15] and on their GitHub repository (https://github.com/broadinstitute/gtex-pipeline/tree/master/qtl/leafcutter). Briefly, reads were first aligned to the reference with STAR (version 2.7.10a)[55], using WASP correction to mitigate allelic mapping bias. For WASP correction, we used phased variant calls from the NYGC's high-coverage sequencing of the 1KGP[23] (20201028 accession, located on the International Genome Sample Resource[56] ftp server here: https://ftp.1000genomes.ebi.ac.uk/vol1/ftp/data_collections/1000G_2504_high_coverage/working/20201028_3202_phased/). All other options were as described by GTEx.

These alignments were then supplied to Leafcutter (version 0.2.9)[16] to generate intron excision ratios. Notably, Leafcutter is annotation agnostic; it defines its own splicing "clusters" (groups of related intron excision events) using split reads rather than quantifying splicing using prior exon or transcript annotations. In a data set such as ours, where many individuals are from historically understudied populations, this eliminates bias from annotations generated from sample sets with limited diversity and may allow us to elucidate novel intron excision events.

Intron usage was estimated for each library using `regtools junctions extract` using a minimum anchor length of 8bp (`-a 8`), strand specificity set to unstranded (`-s 0`) based on our library prep, minimum intron size set to 50bp (`-m 50`), and the maximum intron size set to 500kb (`-M 500000`). Junction files were then used to cluster introns across all samples using the Leafcutter `leafcutter_cluster_regtools.py` companion script, where 50 split reads were required to support each cluster (`-m 50`) and the maximum intron size was set to 500kb (`-l 500000`).

While STAR and Leafcutter were run using all 779 sequencing libraries, unless otherwise stated, for the 24 samples that were sequenced in triplicate, downstream analyses used results from the replicate with the most reads for each of these samples.

#### 5.2 Filtering lowly-expressed and low-complexity clusters

Introns with low counts and clusters with low complexity can lead to statistical issues when discovering sQTLs and quantifying splicing variance. To avoid these issues, we applied a filtering procedure to the intron excision ratios produced by Leafcutter, largely based on the leafcutter filtering applied by GTEx and described in their paper[15] and on their GitHub repository (https://github.com/broadinstitute/gtex-pipeline/blob/master/qtl/leafcutter/src/cluster_prepare_fastqtl.py).

After running Leafcutter, we had intron excision ratios for 245,487 introns (51,466 splicing clusters) on the autosomes and chrX. We first filtered out introns with low complexity across samples, defined as introns without any read counts in >90% of samples, or with fewer than $max(10, 0.1n)$ unique values, where $n = 731$ is the sample size. After this step, 154,816 introns (33,712 clusters) remained. Through use of the Leafcutter `prepare_phenotype_table.py` companion script (described in more detail in **section 11.1** below) 8,430 additional introns were dropped with SD < 0.005 across samples or whose cluster had 0 counts in > 40% of samples. After this step, 146,386 introns (33,447 clusters) remained. Finally, we dropped 580 clusters with only one intron.

After filtering, 145,806 introns (32,867 splicing clusters) were retained and used for down-stream analyses of splicing variation.

### 6 Quantifying the contribution of batch effects to expression variation

Batch effects–along with other sources of technical variation–are a known confounder in RNA-seq data. As such, it is critical to ensure that the effect of batch on gene expression level does not mask actual biological variation. To assess the contribution of batch effects to expression level variation, we sequenced 24 samples in triplicate, as described in the **section 1** above. Using the filtered gene-level counts described in **section 4.2**, we calculated the Spearman rank correlation between each pair of the 72 replicate sequencing libraries. Critically, we observe that pairs of libraries from the same individual have higher correlations than pairs of libraries from the same batch (**Fig. S3A**).

Additionally, for each of the 19,539 autosomal expressed genes and across the 72 replicate sequencing runs (24 samples sequenced in triplicate), we calculated the proportion of expression level variation explained by sample versus batch. Using a VST normalized expression matrix (described in **section 7.1** below) subset to samples sequenced in triplicate, we performed a type II ANOVA using the `Anova` function from the *car* package (version 3.1-2) in R with the following regression formula: *expression ~ batch + sample*. To test whether expression variance between samples was greater than variance measured between sequencing batches, we performed a Wilcoxon signed-rank test using the proportion of variance explained from the ANOVA above. We observed that, on average, the proportion of gene expression variance explained by sample was greater than by sequencing batch ($p < 1 \times 10^{-10}$), concordant with the results from the Spearman's rank correlation test (**Fig. S3B**).

We performed a complementary set of analyses to quantify the contribution of batch effects to splicing variation. Using the filtered intron excision ratios described in **section 5.2**, we calculated the Spearman rank correlation between each pair of the 72 replicate sequencing libraries. As with the analysis of expression level, we observe that pairs of libraries from the same individual have higher correlations than pairs of libraries from the same batch (**Fig. S3C**). Additionally, for each of the 31,837 autosomal splicing clusters that passed filtering, we calculated the proportion of splicing variation explained by sample versus batch. Using the intron excision ratios from Leafcutter, we performed a type II ANOVA using the `manta` function from the *manta* package (version 1.0.0) in R to fit a model that regresses intron excision ratios onto batch and sample. Described in more detail in section **9**, MANTA[57] is a tool for evaluation of multivariate linear models (such as intron excision ratios) that uses the Hellinger distance between splicing ratios to estimate the variability in splicing across individuals. As before, we performed a Wilcoxon signed-rank test using the proportion of variance from MANTA. We observed that, on average, the proportion of splicing variance explained by sample was greater than by sequencing batch ($p < 1 \times 10^{-10}$) - concordant with the results from the Spearman's rank correlation test (**Fig. S3D**).

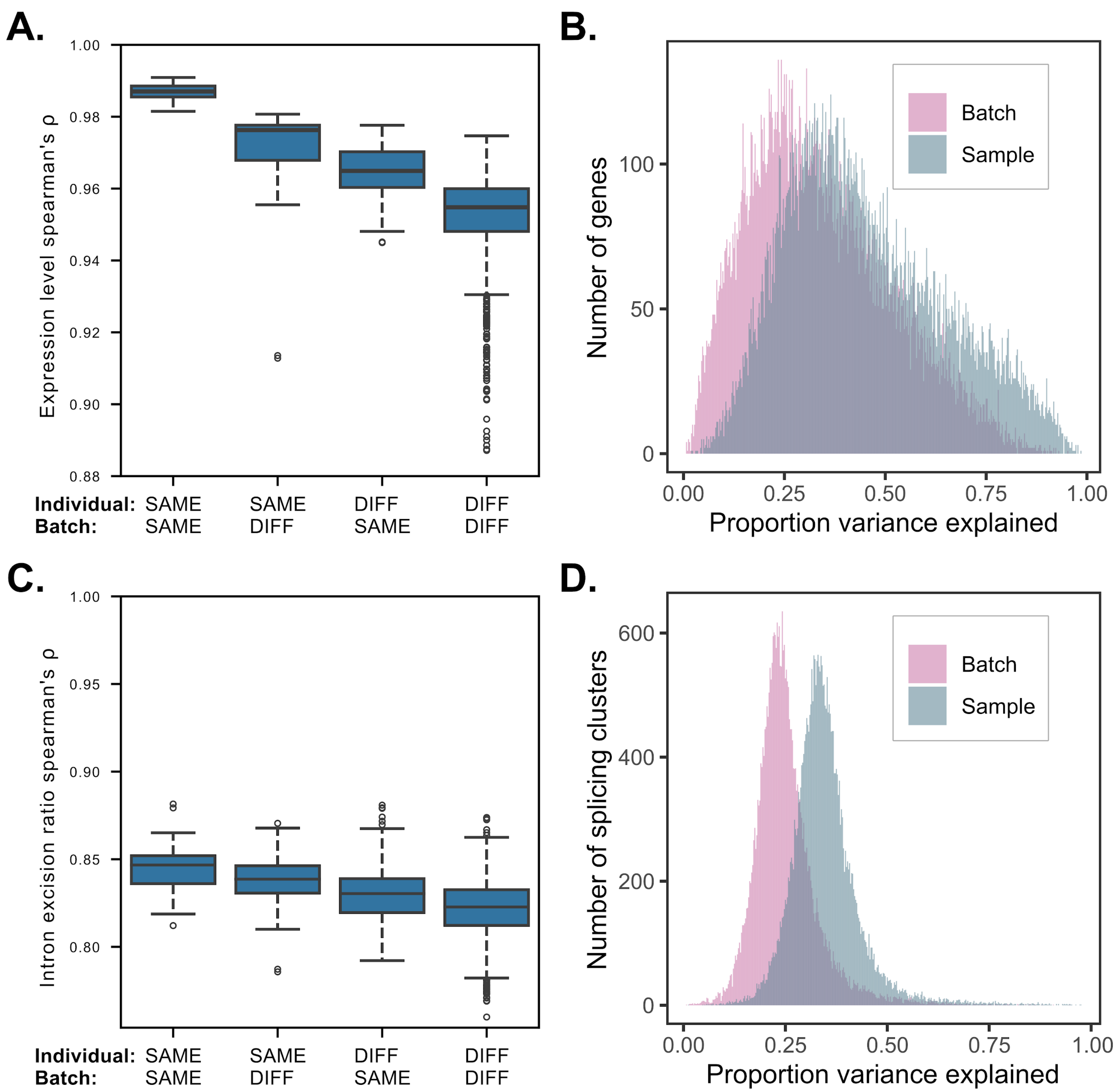


**Figure S3. Batch effects in MAGE RNA-seq data. (A)** Spearman rank correlation in expression level (TMM) across all expression-filtered genes between each pair of technical replicates in our data set (24 unique samples, 72 total replicates). Pairs of replicates are stratified by 1) whether they were sequenced in the same sequencing batch and 2) whether they were derived from the sample 1KGP sample. Higher correlations are observed for pairs of replicates from the same sample than for pairs of replicates from the same sequencing batch. **(B)** For replicate sequencing libraries and across autosomal expression-filtered genes, per gene estimates of the proportion of variance in gene expression level explained by sequencing batch (pink) or sample (blue). On average, sample explained a higher proportion of variance in expression level than batch (Wilcoxon signed-rank test: $p < 1 \times 10^{-10}$). **(C)** Same as panel A, but showing Spearman rank correlation in intron excision ratio across all splicing-filtered introns. Again, higher correlations are observed for pairs of replicates from the same sample than for pairs of replicates from the same sequencing batch. **(D)** Same as panel B, but showing the proportion of variance explained by batch or sample across autosomal splicing-filtered splicing clusters. On average, sample explained a higher proportion of variance in splicing than batch (Wilcoxon signed-rank test: $p < 1 \times 10^{-10}$).

### 7 Differential gene expression between populations

#### 7.1 Data preparation

Differential gene expression (DGE) analysis was performed using *DESeq2* (version 1.36.0[58]) in R. Using the salmon-generated pseudocount expression data per sequence library (detailed in section **4.1**), transcript-level abundances were first converted into gene-level abundances using the *tximport* (version 1.24.0)[54] in R under default parameters. Gene-level expression estimates were imported into the DESeq2 ecosystem with the design formula specified as ~ *population* + *batch* + *sex,* where *batch* and *sex* were included as categorical covariates to control for technical variation between sequencing batches (see section **6**) and sex-dependent effects. For this analysis, technical replicates for each of the 24 samples that were sequenced in triplicate were collapsed into single samples using the `collapseReplicates` function included in DESeq2.

#### 7.2 Factor contrasts

Differential expression contrasts were constructed one of two ways: 1) each population's expression was contrasted against the average expression across all other populations within their parent continental group (e.g., JPT samples vs. all other samples within the East Asian continental group), or 2) each continental group's expression was contrasted against the average expression across all other continental groups (e.g., AFR samples vs. all other samples).

Using the design formula specified in **section 7.1**, Wald test contrast coefficient matrices were extracted for each population by computing the mean coefficients for each dummy variable using all samples within the focal population label (e.g., JPT). For each continental group and background population (e.g., all non-focal subpopulations per continental group), coefficient matrices were additively combined and normalized to the number of populations contained within their respective continental group (yielding a contrast coefficient matrix where the intercept weight = 1). Multiple testing correction, independent filtering, and outlier detection for each contrast were all performed using default functions included in the *DESeq2* package.

### 8 Expression level variation within and between populations

#### 8.1 Normalized expression matrix

The relationship between populations and expression variance was measured using the blind variance stabilizing transformation (VST) function included in *DESeq2* (version 1.36.0)[58] on the gene-level count matrix produced after collapsing technical replicates (see section **7.1**). VST produces an expression matrix which directly captures the effects of library- and experiment-wide normalization factors, estimated gene-wise dispersion, and reduces the dependence of expression variance on the mean expression per-gene (see **Fig. S4**). This transformed count matrix was reduced to only represent the filtered subset of genes described in section **4.2**.

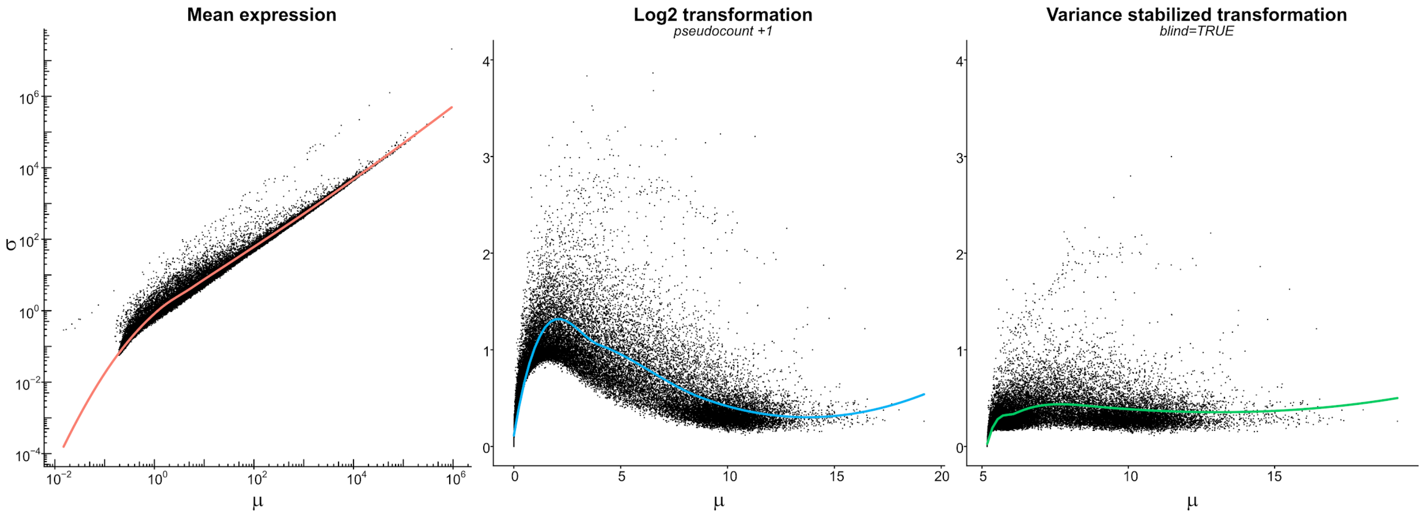


**Figure S4. Count-data transformations for examining global trends in gene expression.** Experiment-wide expression mean, μ, and standard deviation, σ, computed across all samples following three methods of expression normalization: mean expression computed by DESeq2 after correcting for dispersion and normalization factors (left), $Log_2$-transformed counts (middle), and variance-stabilized transformation (VST; right).

#### 8.2 Estimation of biological variation

We applied a two-stage ANOVA strategy to quantify gene expression variance at global, continental group, and population scales. First, an ANOVA was performed for each gene using the `anova` and `lm` functions in the *stats* package (version 4.3.0) in R with formula **[1]** below, where *batch* and *sex* were included as categorical covariates to remove technical variance from the response variable. Here *u* is the residuals of the regression, and represents the VST normalized expression values corrected for the effects of batch and sex. Because continental group and population together form a multicollinear system, two independent ANOVAs were then performed to estimate the proportion of gene expression variance due to continental group (formula **[2]**) and population label (formula **[3]**), where the batch- and sex-corrected expression values, *u*, were used as the response variable. The proportion of variance explained (PVE) was estimated as the regression sum of squares divided by the total sum of squares for each regression. In this manner, the PVE by continental group or by population represent the proportion of variance in the batch- and sex-corrected expression values explained by the label.

$$\textbf{(1)}\quad VST\ Expression \sim sex + batch\ + u$$

$$\textbf{(2)}\quad u \sim continental\ group\ +\ v_1$$

$$\textit{(3)}\quad u \sim population\ +\ v_2$$

To test whether the variance explained by continental group/population was greater than that expected by chance, we performed a permutation test where continental group and population labels were shuffled (with replacement) and the ANOVA procedure described above was recalculated for each gene (n = 1000 permutation replicates). A permutation test p-value was computed as the proportion of permutations where the mean proportion of variance explained by continental group/population was more extreme (i.e., greater) than those respectively measured in our empirical data set. For both continental group and population, none of the permutations had a mean PVE greater than calculated with the empirical data set.

To quantify variance in gene expression within each continental group, we first applied the same regression strategy to remove variance due to sex and batch from the VST gene expression array using formula **[1]**. For each continental group, and for each gene, residuals were partitioned to include only samples within the focal group, and sample variance was calculated using the `var` function in R.

Using the gene-wise variance estimates per continental group, we tested whether gene expression differs significantly across continental populations (**Fig. S5A**). To achieve this objective, we fit a linear mixed model (`lme`, *lme4* package version 1.1-34 in R) to the expression data, where the response variable ($log_{10}$-transformed variance) was regressed against a continental group fixed-effect and gene included as a random-effect. The performance of this model was compared with a reduced model (without the continental group fixed effect) using the `anova` function in the *stats* package. This statistical procedure was also applied to test whether splicing differs significantly across continental groups (**Fig. S5B**).

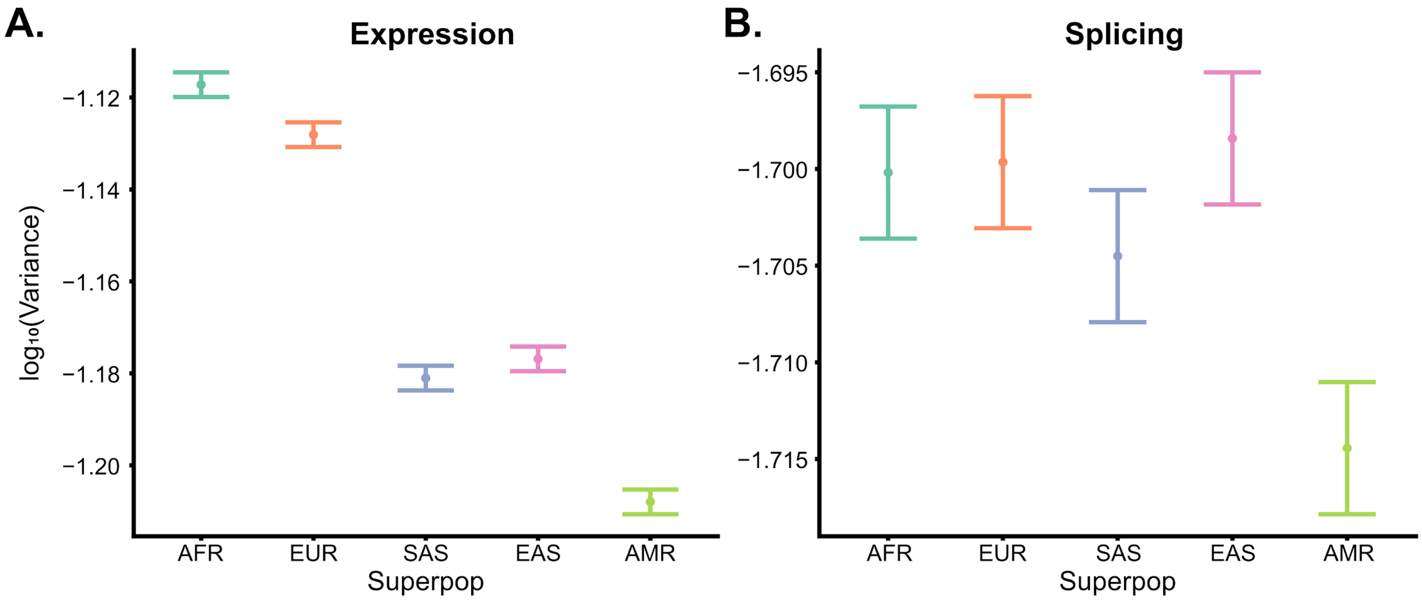


**Figure S5. Global trends in gene expression and splicing variance. (A)** Variance in gene expression variance decreases with Eastward expansion from Africa. **(B)** In contrast, splicing variance shows little change with Eastward expansion, decreasing significantly (ANOVA p-value = 3.17e-3) in the admixed American continental group (relative to the African continental group). Each point represents the mean variance computed across all genes/splicing clusters within the focal population label, and whiskers represent ± 1 std. Err.

### 9 Splicing variation within and between populations

The approach to quantify splicing variation between and within continental groups and populations largely mirrors the approach used for gene expression level (detailed in section **8.2**).

As with gene expression level, we applied a two-stage ANOVA strategy to quantify the proportion of splicing variance explained by continental group or population. Because splicing proportions are inherently multivariate, a standard ANOVA is not appropriate. Instead, we used MANTA[57], a tool for evaluation of multivariate linear models including proportion data such as intron excision ratios. MANTA uses the Hellinger distance between splicing ratios to estimate the variability in splicing across individuals.

For each splicing cluster that passed filtering (see section **5.2**), we applied the following procedures. First we used the `manta` function from the *manta* package (version 1.0.0) in R to regress filtered intron excision ratios onto sample-batch and sex to remove technical variation from the response variable. We set `transform="sqrt"` to use the Hellinger distance between splicing ratios, and `fit=TRUE` to return the regression residuals. Using the residuals from this first step, we then ran two independent ANOVAs to estimate the proportion of splicing variance attributable to continental group and population label. As before, we used the `manta` function to regress the residuals from the first step onto either continental group or population. We did not use the square root transform for these two models, because the residuals from the first step should reflect the initial square root transform. The proportion of variance explained by either continental group or population was estimated as the regression sum of squares divided by the total sum of squares (after regressing out batch and sex) for each model.

As with gene expression level, we tested whether the variance explained by continental group/population was greater than that expected by chance using a permutation test. Continental group and population labels were shuffled and the above procedure was repeated. 1000 total permutations were performed. We computed a permutation test p-value for both continental group and population as the proportion of permutations where the mean PVE (across splicing clusters) was greater than that calculated from the empirical data set. For both continental group and population, none of the permutations had a mean PVE greater than calculated with the empirical data set.

To quantify variance in splicing within each continental group, we first applied the same regression strategy described above to remove variance from batch and sex from intron excision ratios. For each continental group, and for each splicing cluster, residuals from this regression were partitioned to include only samples within the focal continental group, and sample variance was calculated as:

$$\frac{1}{N(N-1)} \sum_{j}^{N-1} \sum_{k=j+1}^{N} d^2(j,k)$$

where $N$ is the total number of samples within the focal continental group, and $d^2(j,k)$ is the squared Euclidean distance between the residual intron excision ratios (after removing the effects of batch and sex) of the focal splicing cluster for individuals $j$ and $k$ in the focal continental group.

### 10 *cis*-eQTL mapping

#### 10.1 Expression normalization

Preliminary gene expression counts (described in section **4.1**) were prepared for eQTL mapping using the following procedure: 1) gene-level counts were normalized between samples using TMM[59] as implemented in the *EdgeR* package (version 3.32.1)[60] in R; 2) lowly expressed genes were filtered out (as described in section **4.3**); 3) for each gene that passed filtering, TMM values were inverse normal transformed.

#### 10.2 Calculation of genotype PCs

To control for the effects of global ancestry on gene expression, we first calculated the top 20 genotype principal components (PCs) from the samples included in MAGE. Genotype PCs were computed from the NYGC high-coverage variant calls (see section **2**) across the 731 samples using PLINK[51] with the `--pca` option and restricting to variants with in-sample MAF > 0.01 using `--maf 0.01`. We observed that the variance explained by consecutive PCs decreased considerably following the first five genotype PCs (**Fig. S6A**). Additionally, the top four genotype PCs correlated strongly with continental group label (**Fig. S6B**), and the top five genotype PCs correlated strongly with population label (**Fig.**

**S6C**). Interestingly, we do observe some weaker correlations with population label; for example, PC10 appears to be correlated with population label, but explains only 0.26% of genotype variance.

Based on these results, the top five genotype PCs were included as covariates in QTL mapping to control for confounding by global ancestry.

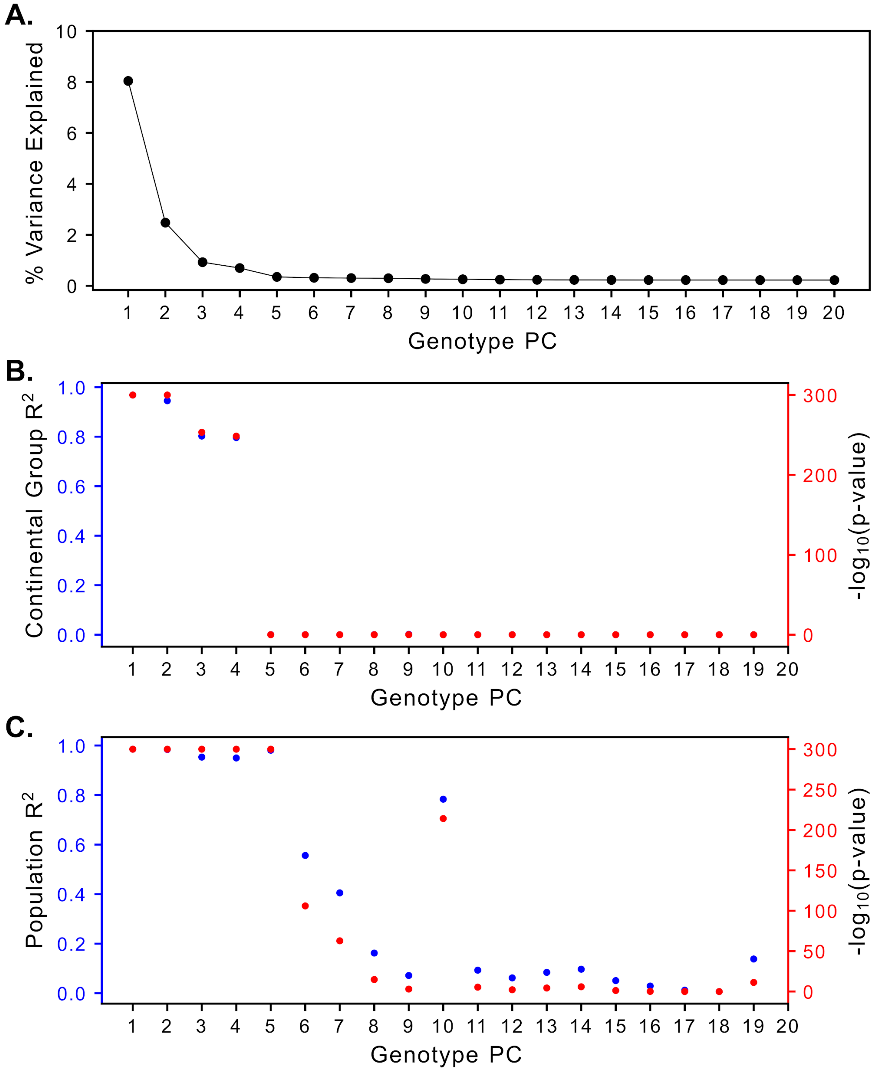


**Figure S6. Selection of genotype PCs for QTL mapping. (A)** Percent of genotype variance explained by each of the top 20 genotype PCs for the samples in MAGE. Variance explained drops off after 5 PCs. **(B)** Correlation between continental group and each of the top 20 genotype PCs. The first 4 PCs are significantly correlated with sample continental group label. **(C)** Correlation between population label and each of the top 20 genotype PCs. The first 5 PCs are significantly correlated with population label. Interestingly, PC10 also appears to be correlated with population label, but explains only 0.26% of genotype variance.

#### 10.3 Calculation of PEER covariates

Batch effects and other technical sources of variation are known to affect RNAseq studies and quantification of gene expression, and can reduce the power of eQTL mapping if not properly controlled. Because these factors are not necessarily directly measured, we used Probabilistic Estimation of Expression Residuals (PEER)[61] to identify hidden factors driving expression variation in our data set. Based on the optimizations performed previously by GTEx[15,61], we calculated 60 PEER factors to use as covariates in eQTL mapping. We used `peertool` (v1.0) to calculate 60 PEER covariates from the normalized TMM values (see section **10.1**), limiting the algorithm to 100 iterations using `--n_iter 100`.

#### 10.4 Discovery of nominal *cis*-eQTLs with FastQTL

We discovered eQTLs using FastQTL (version v2.184_gtex)[62] as implemented by GTEx (https://github.com/francois-a/fastqtl). For each of the 19,539 autosomal genes that passed filtering thresholds (see section **4.2**), we regressed inverse normal transformed TMM values onto variant genotypes for all variants within 1 Mbp up- and down-stream of the gene's transcription start site (TSS), and with MAF > 0.01. The top 5 genotype PCs (section **10.2**), 60 PEER factors (section **10.3**), and sex were included as covariates.

We first ran FastQTL in the adaptive permutation mode using `--permute 1000 10000`, to discover significant *cis*-eGenes (genes with at least one cis-eQTL). FastQTL estimates gene-level empirical p-values, based on the theoretical distribution of permutation p-values. The GTEx implementation of FastQTL uses the estimated empirical p-values to calculate gene-level q-values and from these q-values, we discover eGenes at a 5% false discovery rate (FDR) threshold. FastQTL also calculates a nominal p-value threshold for significance for each gene, based on the chosen FDR.

To identify significant *cis*-eQTL associations, we ran FastQTL in a nominal pass (the default), and defined significant *cis*-eQTLs as those variant-gene pairs whose nominal p-value was below the nominal p-value threshold (at a 5% FDR) for the tested gene.

#### 10.5 Fine-mapping eGene credible sets with SuSiE

To discover the causal SNP(s) driving each *cis*-eQTL signal, we performed fine-mapping with SuSiE[24,63], using the *susieR* package (version 0.12.16) in R. For each tested gene, SuSiE discovers a set of credible causal sets, such that each has some minimum probability of containing a true causal SNP (termed the "coverage" probability), each credible set is made as small as possible, and SNPs within each credible set have some minimum correlation with each other. As such, SuSiE can discover multiple independent signals per gene, and at high resolution.

For each *cis*-eGene identified at a 5% FDR with FastQTL (see section **10.4**), we identified SuSiE credible sets using the following procedure. For each gene, we limit the analysis to the same set of SNPs used with FastQTL, specifically SNPs within 1Mbp up- and down-stream of the gene's TSS, and with MAF > 0.01. We then remove the effects of the eQTL-mapping covariates (sex, top 5 genotype PCs, 60 PEER factors) from the inverse normal transformed TMM values and genotypes, using the procedure described in this article: https://stephenslab.github.io/susieR/articles/finemapping.html#a-note-on-covariate-adjustment. Finally, we run the `susie_rss` function on the Z-scores from the FastQTL nominal pass, using an in-sample LD matrix calculated from the covariate adjusted genotypes and gene expression variance estimated from the covariate-adjusted expression values. We set the maximum number of credible sets to be 10 (`L=10`), the minimum coverage probability of each credible set to be 0.95 (`coverage=0.95`), and the minimum absolute correlation between SNPs in a credible set to 0.5 (`min_abs_corr=0.5`).

SuSiE discovered credible sets for 9,807 of the 15,022 eGenes identified in the FastQTL permutation pass. For each fine-mapped credible set, we select a single representative "lead" eQTL with the highest PIP within that credible set. We use these lead eQTLs in all downstream analyses to represent putative causal eQTL signals.

#### 10.6 Comparison of fine-mapping resolution in subsets of MAGE

To investigate the relationship between fine-mapping resolution and sample diversity of the discovery set, we repeated our eQTL-mapping pipeline for three equally sized (n = 142) subsets of the MAGE data set: one that included only samples in the AFR continental group of 1KGP, a second that included only samples in the EUR continental group of 1KGP, and a third that included samples from all 26 populations of 1KGP. Within each subset, samples were selected from the populations included in the subset as evenly as possible. For each subset, we independently repeated the entire eQTL mapping pipeline (sections **10.1-10.5**) as before with two minor changes, both related to the smaller size of the sample: 1) the MAF cutoff for eQTL mapping was set to 0.05 rather than 0.01 and 2) only 15 PEER factors were calculated and included instead of 60.

We compared the size of the resulting SuSiE credible sets 1) for genes with at least one SuSiE credible set in any of the subsets (**Fig. S7A**), and 2) for genes with at least one SuSiE credible set in all three subsets (**Fig. S7B**). In both cases, we observe the best resolution (fewest variants per credible set) on average in the African subset, the second-best resolution in the diverse subset, and the worst resolution in the European subset. This result is expected given the increased genetic diversity in African populations[21,22], and highlights the advantages that inclusion of diverse samples affords for detection of causal signals.

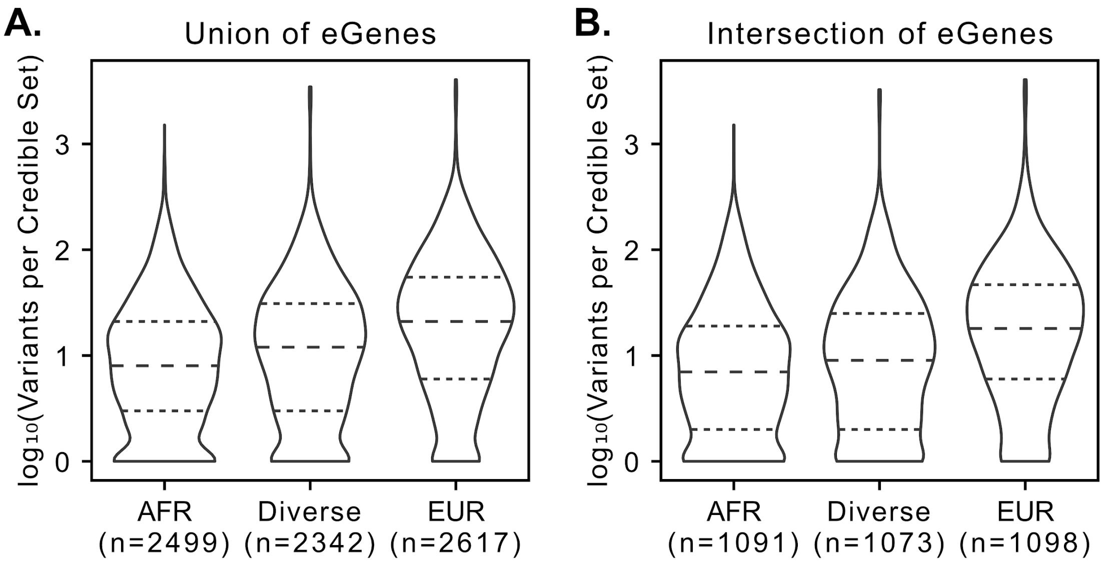


**Figure S7. Comparison of fine-mapping resolution in subsets of MAGE.** We re-ran the entire eQTL mapping pipeline (including filtering of variants and genes) for three equally sized (n = 142) subsets of the MAGE data set, one that included only samples in the AFR continental group of 1KGP, a second diverse subset that included samples from all 26 populations of 1KGP, and a third that included only samples in the EUR continental group of 1KGP. The pipeline was run separately for each subset. **(A)** The number of variants per SuSiE credible set within each subset for all genes that had at least one credible set in at least one subset. **(B)** Same as panel A, but only for those genes that had at least one credible set in all three subsets. The AFR subset yields the smallest (best resolution) credible sets, as expected given the increased genetic diversity in African populations[21,22]. Importantly, the diverse subset yields only slightly lower resolution credible sets. The worst resolution is observed in the EUR subset.

#### 10.7 Calculation of Allelic Fold Change (aFC)

While valuable for identifying significant eQTLs, the slope of the regression from the FastQTL nominal pass (see section **10.4**) is based on rank-normalized expression quantifications and as such does not have a clear biological interpretation. At the same time, for eGenes with multiple independent causal signals, the measured "nominal" effect size of one causal eQTL can be influenced by the effects of the other causal signals. As such, for each eGene, it is useful to calculate the "marginal" effect size of each causal signal, conditional on the effects of the other causal signals for that gene. This better reflects the actual effect each causal signal has on the expression of its eGene.

One metric for quantifying the effect of an eQTL on the expression of its eGene is allelic fold change (aFC), which describes the ratio between the expression of the haplotype carrying the alternative allele to the one carrying the reference allele[26]. This concept can be extended to handle multiple causal signals per gene, as implemented in the aFC-n tool[64]. For each lead eQTL in our fine-mapping results (see section **10.5**), we calculated the effect size of that variant as log2(aFC) using the following procedure:

First, we generated corrected expression counts for each gene using using DESeq2 (version 1.36.0; [58]) in R. Using the salmon-generated pseudocount expression data per sample (detailed in section 4.1), transcript-level counts were first converted into gene-level counts using the tximport (version 1.24.0; [54]) in R under default parameters. These gene-level expression estimates were imported into the DESeq2 ecosystem, and we generated "corrected" expression counts, using the `counts` function with `normalized=TRUE`. These corrected counts are functionally equivalent to read counts but have been corrected for library size and average transcript length. These corrected counts were then log2-transformed (with a +1 pseudocount).

Next, we removed the effects of covariates using the following procedure: for each eGene we fit a linear model that regresses $\log_2$(corrected counts) onto sample genotypes for each lead eQTL of that gene as well as the eQTL-mapping covariates described in section **10.4**. Any covariates whose 95% confidence interval did not include 0 were regressed out from the $\log_2$(corrected counts).

Finally, we used `afcn.py` with the `--conf` option, using these covariate-adjusted $\log_2$(corrected counts) as input, to calculate $\log_2$(aFC) for each lead eQTL (**Fig. S8**).

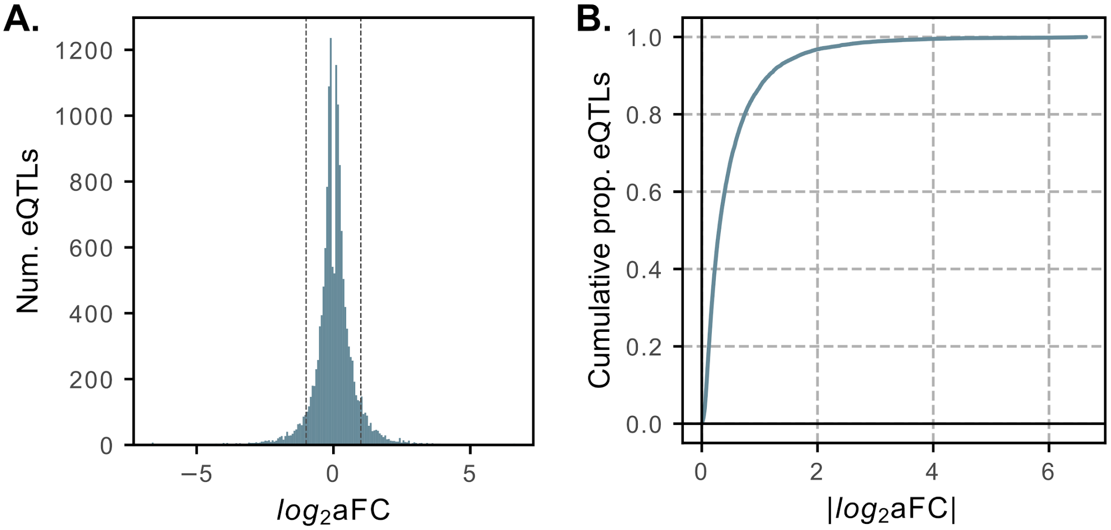

**Figure S8. Distribution of eQTL effect sizes. (A)** Distribution of lead eQTL effect sizes, measured as $\log_2(\text{aFC})$. This distribution is expected to be roughly symmetric as, for each variant, the sign of the effect is entirely dependent on which allele is denoted the reference allele. Vertical dotted lines denote a two-fold change to expression ($\log_2(\text{aFC}) = \pm 1$). Most eQTLs have a relatively small effect on expression level. **(B)** Cumulative distribution of the absolute value of effect size across lead eQTLs. Only 2031 (13%) lead eQTLs had greater than a twofold effect on gene expression (median $|\log_2(\text{aFC})| = 0.30$).

### 11 *cis*-sQTL mapping

#### 11.1 Splicing normalization

Intron-excision ratios from Leafcutter were filtered as described in section **5.2**. Prior to removing splicing clusters with a single intron (but after removing low complexity introns), intron excision ratios were normalized using the Leafcutter `prepare_phenotype_table.py` companion script. We then filtered out splicing clusters with only a single intron. Splicing clusters were mapped to annotated genes in GENCODE v38[52] using the Leafcutter `map_clusters_to_genes.R` companion script. Of the 32,867 splicing clusters remaining after filtering, we removed 679 additional clusters that did not map to annotated exons in GENCODE v38.

For sQTL mapping, normalized intron excision ratios (across all samples) were collected into a bed file, with each cluster annotated with the TSS of the gene to which it mapped. If a cluster mapped to multiple genes, each mapping was included in the bed file separately.

#### 11.2 Calculation of PEER covariates

For sQTL mapping, PEER factors were calculated from the normalized intron excision ratios described in section **11.1**. Based on the optimizations performed previously by GTEx[15], we calculated 15 PEER factors to use as covariates in sQTL mapping. Otherwise, PEER factors were calculated as described in section **10.3**.

#### 11.3 Discovery of nominal *cis*-sQTLs with FastQTL

The *cis*-sQTL mapping procedure largely matched the procedure used to map *cis*-eQTLs, as described in section **10.4**. For each of the 11,912 autosomal genes with splicing clusters that passed filtering thresholds (see section **11.1**), we regressed normalized intron excision ratios onto variant genotypes for all variants within 1Mbp up- and down-stream of the gene's transcription start site (TSS), and with MAF > 0.01. The top 5 genotype PCs (section **10.2**), 15 PEER factors (section **11.2**), and sample sex were included as covariates.

As with *cis*-eQTL mapping, we first ran FastQTL in the adaptive permutation mode using `--permute 1000 10000` to discover significant *cis*-sGenes (genes with at least one *cis*-sQTL).We used grouped permutations (using the `--phenotype_groups` option) to compute a gene-level empirical p-value over all splicing clusters of a gene. We discovered *cis*-sGenes and calculated per-gene nominal p-value thresholds at a 5% FDR (as described in section **10.4**).

To identify significant *cis*-sQTL associations, we ran FastQTL in a nominal pass and defined significant *cis*-sQTLs as those variant-intron pairs whose nominal p-value was below the 5% FDR nominal p-value threshold for the tested gene.

#### 11.4 Fine-mapping sGene credible sets with SuSiE

For each 5% FDR *cis*-sGene identified, fine-mapping was performed separately for each intron mapping to that gene (termed sIntrons; limited to only the introns that passed filtering). Fine-mapping was done as described for *cis*-eGenes (section **10.5**), mapping normalized intron excision ratios onto sample genotypes, using the same set of covariates used for sQTL mapping with FastQTL. All other options remained the same.

SuSiE discovered credible sets (for at least one intron) for 6,604 of the 7,727 sGenes identified in the FastQTL permutation pass. Of the 25,864 fine-mapped sIntrons 4425 (17%) had more than one credible set (**Fig. S9A**), representing 1,777 (27%) of the 6,604 fine-mapped sGenes. Of the 32,436 intron-level credible sets, 7,718 (24%) contained just a single variant (median 6 variants per credible set; **Fig. S9B**).

To obtain a gene-level summary of the sQTL fine-mapping results, we collapsed these intron-level credible sets into gene-level credible sets. For each sGene, we iteratively merged all intron-level credible sets that overlapped by at least one variant. The result is a set of gene-level merged credible sets that are independent from one another (no variants in common) and whose union is equivalent to the union of the input intron-level credible sets. Of the 6,604 fine-mapped sGenes, 3,490 (53%) had more than one credible set (**Fig. S9C**). Of the 16,451 gene-level credible sets, 3,569 (22%) contained just a single variant (median 7 variants per credible set; **Fig. S9D**).

Analogous to selection of lead eQTLs described in section **10.5**, we selected a representative "lead" sQTL for each gene-level merged credible set by first determining the intron-level credible set with the greatest coverage among those comprising the gene-level merged credible set, and then selecting the variant within that intron-level credible set with the highest PIP. We use these lead sQTLs in all downstream analyses to represent putative causal sQTL signals.

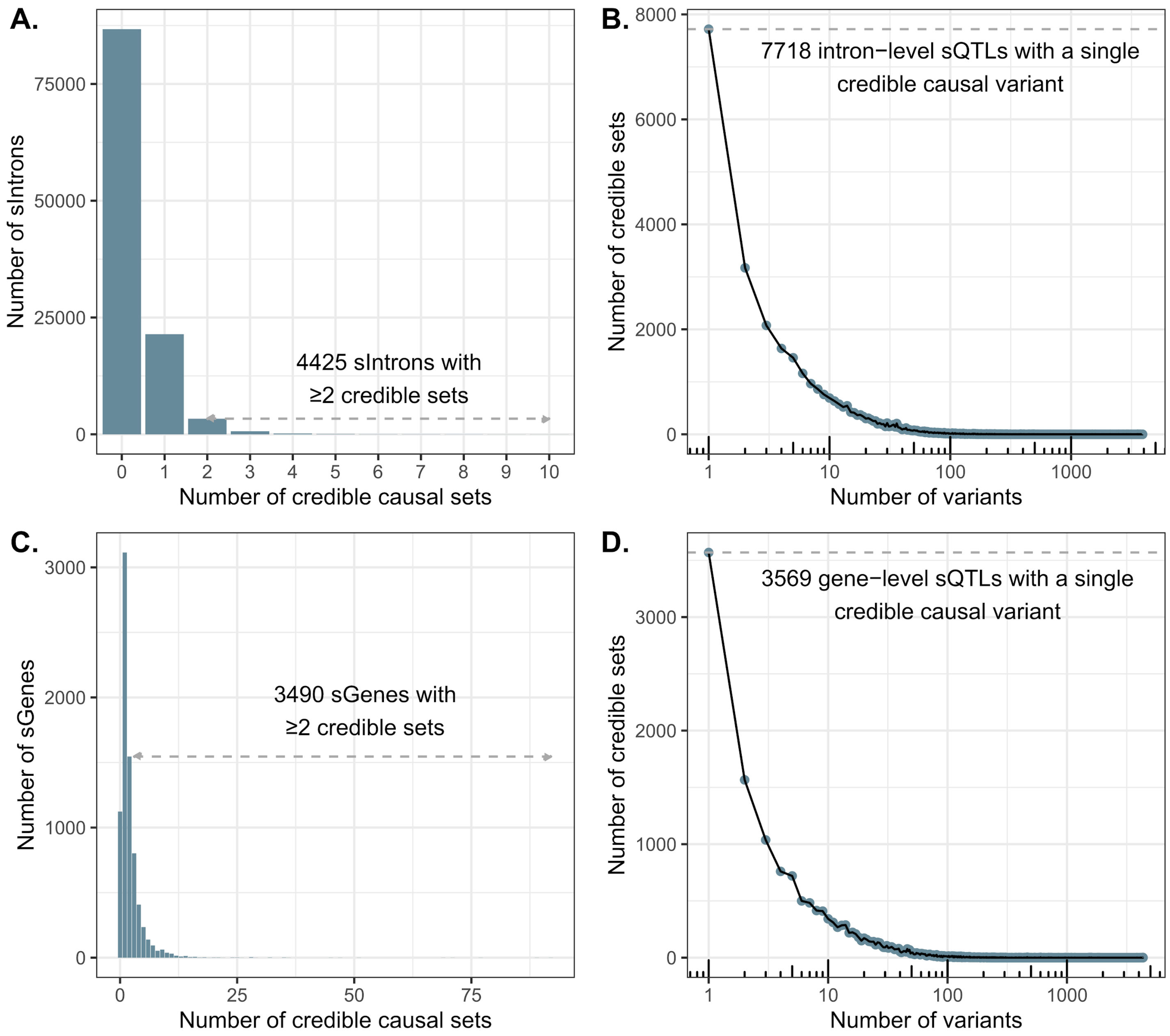


**Figure S9. Mapping of high-resolution sQTLs. (A)** Number of credible sets per sIntron, where we define sIntrons as all introns (that passed filtering) for genes identified as sGenes in the FastQTL permutation pass. We ran SuSiE separately for each sIntron. **(B)** Resolution of sIntron fine-mapping, defined as the number of variants per credible set. **(C)** After fine-mapping, overlapping intron-level credible sets were iteratively merged to produce gene-level credible sets. Panel C shows the number of merged credible sets per sGene. **(D)** The resolution of sGene fine-mapping, defined as the number of variants per merged credible set. These results demonstrate evidence of widespread allelic heterogeneity whereby multiple causal variants independently modulate splicing patterns of the same genes.

### 12 Analysis of negative selection

Evidence of negative selection on regulatory variation affecting highly constrained genes was assessed by intersecting our eQTL fine-mapping results with metrics of constraint generated in previous studies based on depletion of loss of function point mutations or copy number variation[28,65–68] (**Fig. S10**). We restricted our analysis to autosomal genes exceeding the minimum expression threshold used in our differential expression and eQTL mapping pipelines. Among this set of genes, we identified the top 10th percentile of highly constrained genes based on each constraint metric, accounting for their differences in directionality (e.g., high pLI scores but low RVIS scores denote evidence of constraint). The remaining 90% of genes were considered as the background for comparison. We then contrasted the number of independent credible causal sets per gene between the constrained and background set using a quasi-Poisson generalized linear model where normalized mean expression level (i.e., `baseMean`, as computed with DESeq2[58]) was included as a continuous numerical covariate. We also compared the distributions of effect sizes of the lead variant per credible causal set between constrained and background sets of genes using a Mann–Whitney U test, where the base-2 logarithm of the the absolute value of the estimated allelic fold change ($|\log_2(aFC)|$) was used as input to each model, as computed with aFC-n[64].

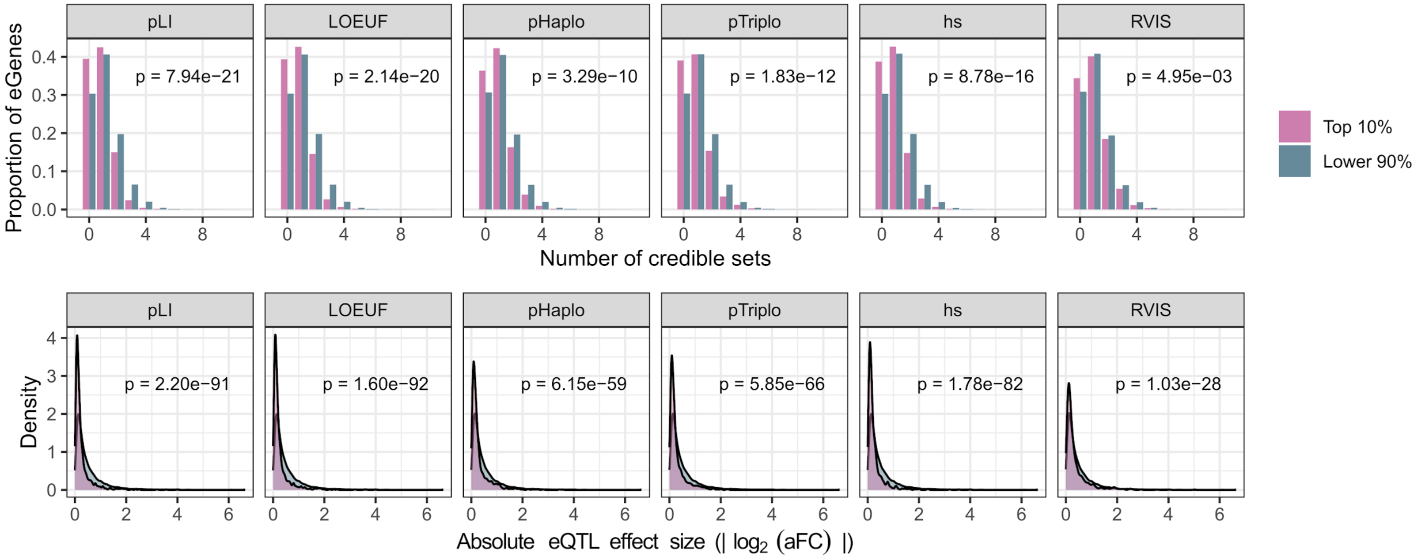


**Figure S10. Evidence of stabilizing selection on gene expression across a range of mutational constraint metrics.** Top row: number of credible causal sets for genes in (pink) and outside (blue) the top decile of various constraint metrics (pLI[66], LOEUF[66], pHaplo[67], pTriplo[67], hs[68], RVIS[65]) obtained from the literature. Bottom row: effect sizes (|log2(aFC)|) of lead eQTLs within (pink) and outside (blue) the same categories.

### 13 Functional annotation and enrichment of fine-mapped *cis*-QTLs

We performed a functional enrichment analysis in order to evaluate the association between fine-mapped *cis*-eQTLs and transcription factor (TF) as well as chromatin regulator (CR) binding sites. The data for this analysis was obtained from the ENCODE Project Consortium, specifically the ENCODE regulation track transcription factor binding site cluster ChIP-seq index file, which encompasses information for 338 DNA-binding proteins across 129 cell types[31,69]. Specifically, we employed GenomicsRanges[70] and BEDTools version 2.29.2 [71] to intersect *cis*-eQTL variants with TF binding sites. Our subsequent enrichment tests were performed using the GREGOR Perl-based pipeline[72]. At a high level, this involves summing the binomial random variables corresponding to the count of index SNPs located within any given TF feature, followed by the computation of enrichment p-values via saddlepoint approximation.

We defined the criterion for positional overlap between SNPs and regulatory features as a minimum of one base pair (≥1 bp) intersection. The fold enrichment for each transcription factor binding site was then calculated as a ratio, defined as the observed fraction of index SNPs overlapping the TF binding sites divided by the expected mean overlap with a matched control SNP set. The control SNPs were matched by the index SNPs' minor allele frequencies and their proximity to the nearest gene's transcription start site (TSS distance), thereby providing a robust basis for comparison ensuring that any observed enrichment is not due to underlying biases in SNP distribution with respect to allele frequency or genomic location. Statistical significance for enrichment was assessed against a background distribution of matched control SNPs, with Bonferroni correction for multiple hypothesis testing to control the family-wise error rate.

To assess chromatin association of the lead eQTLs, we quantified the enrichment of lead eQTLs within the core 15-predicted chromatin states from the Roadmap Epigenomics Consortium[30], which was produced using ChromHMM v1.10 [73], based on a multivariate hidden Markov model. The model delineates the genome into 15 distinct chromatin states based on the combined presence of five key histone modifications: H3K4me3, H3K4me1, H3K36me3, H3K27me3, and H3K9me3. To evaluate enrichment, we examined all 127 reference epigenomes from the Roadmap Epigenomics Consortium encompassing diverse cell types and tissues to ensure a broad representation of epigenetic landscapes, for which we utilized consolidated narrowPeak files for each of 127 epigenomic mappings (**Fig. S11A**). To further parse cell type-specific patterns and consider the predicted enrichment across cell/tissue types, we quantified the enrichment in primary DNAse Hypersensitivity Sites (DHS) data across a diverse panel of 53 cell and tissue types provided by the Roadmap Epigenomics Consortium (**Fig. S11B**).

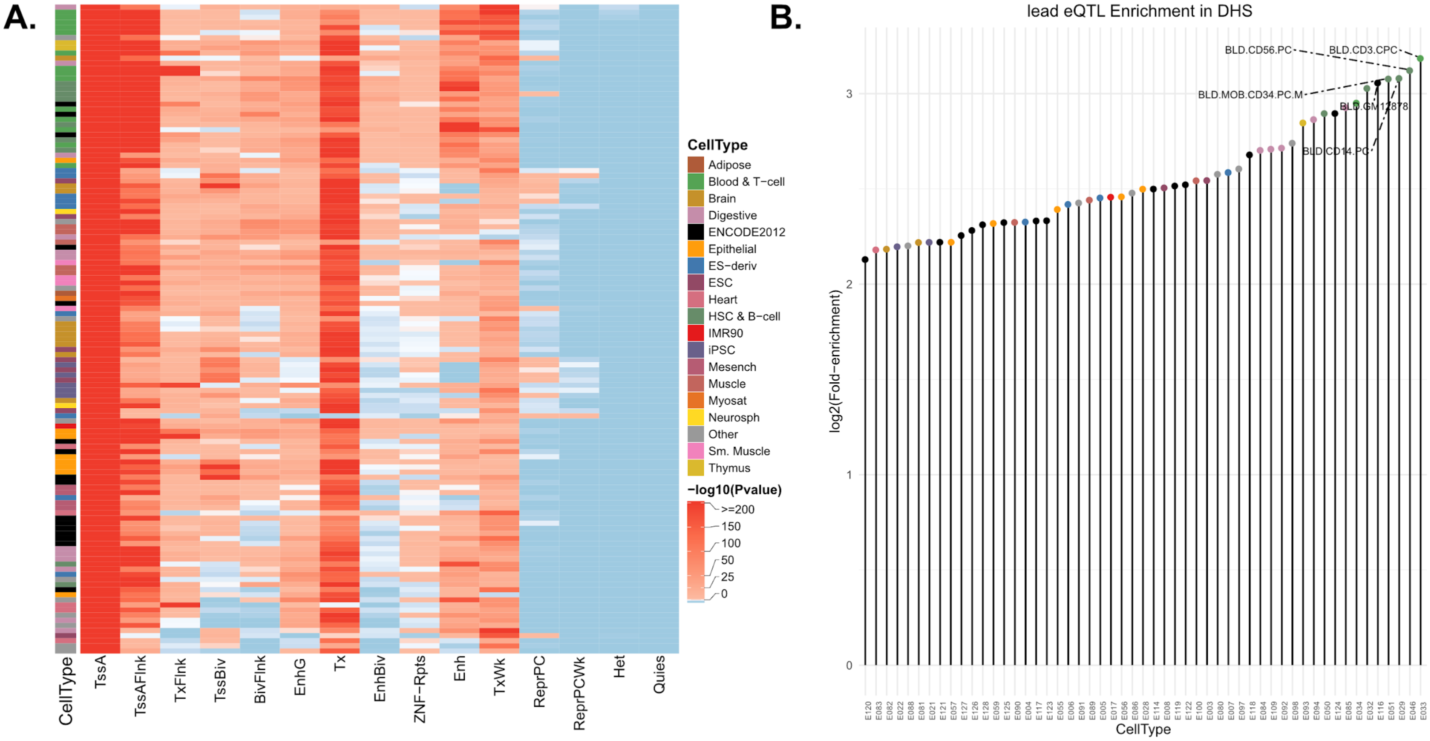


**Figure S11. Lead eQTLs are strongly enriched within regulatory domains across various cell and tissue types. (A)** Corresponding heatmap to **Fig. 4A**, showing significance of the enrichment estimates (right tailed, Binomial P-value). Differential eQTL enrichment across various chromatin states in multiple cell types, highlighting pronounced enrichment at active transcription start sites (TssA) and proximal flanking regions (TssAFlnk), with moderate enrichment in enhancer regions (Enh, EnhG), particularly in blood cell types. Contrastingly, regions characterized by quiescence, repression, and heterochromatin show a marked depletion of eQTLs. **(B)** A lollipop plot showing the pronounced enrichment of lead eQTLs in the DHSs (DNAse Hypersensitivity Sites) across 53 cell/tissue types (colored as in A). We note a marked enrichment in DHS of blood cell types with lymphoblastoid cell line GM12878 as one of the top hits.

We also assessed the distribution of eQTL effect sizes across all 15 predicted chromatin states as annotated with ChromHMM by the Roadmap Epigenomics Consortium. The effect sizes were quantified as the base-2 logarithm of the the absolute value of the estimated allelic fold change ($|\log_2(\text{aFC})|$) across 15 different chromatin states specific to LCLs. Next, to elucidate the regulatory potential of lead *cis*-eQTLs, we assessed the distribution of their effect sizes across promoter, enhancer, and dyadic regions in LCLs associated to multi-tissue DHS data to ensure a comprehensive evaluation of active chromatin domains. Median cis-eQTL effect sizes were compared across these regions to discern any preferential associations. We further stratified eQTLs by effect size, delineated into deciles of absolute $\log_2$ allelic fold change ($|\log_2(\text{aFC})|$), and analyzed their enrichment within the chromatin states predicted for LCLs, including other primary blood cell types. Critically, we also examined how these patterns generalize to other primary blood cell types, including Primary B-cells, T-cells, Natural Killer Cells, and Hematopoietic Stem Cells (**Fig. S12-S16**).

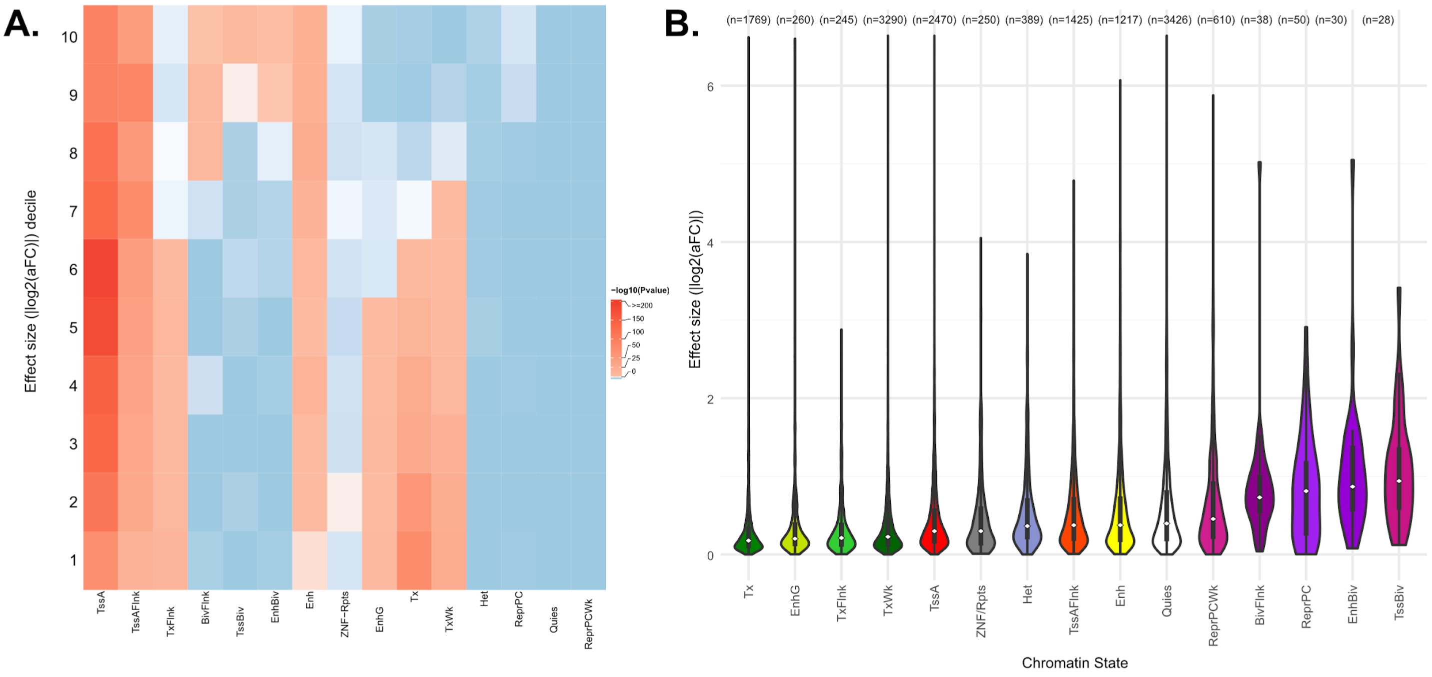


**Figure S12. Lead eQTLs are functionally enriched in the regulatory regions of lymphoblastoid cell lines GM12878 (E116). (A)** Corresponding heatmap to main **Fig. 4B**, showing significance of the decile-based enrichment analysis estimates (right tailed, Binomial P-value). Figure illustrates a heatmap of eQTL effect size distribution, showing consistent promoter-associated enrichment across deciles. Conversely, significant enrichment peaks in bivalent regions (TssBiv, EnhBiv, BivFlnk) are distinctly observed among eQTLs with the largest effect sizes. **(B)** Distribution of effect sizes for lead eQTLs within Roadmap Epigenomics chromHMM predicted chromatin states[30] exhibiting pronounced trend of diminished effect sizes for transcriptional elongation regions (Tx, TxWk, and TxFlnk).

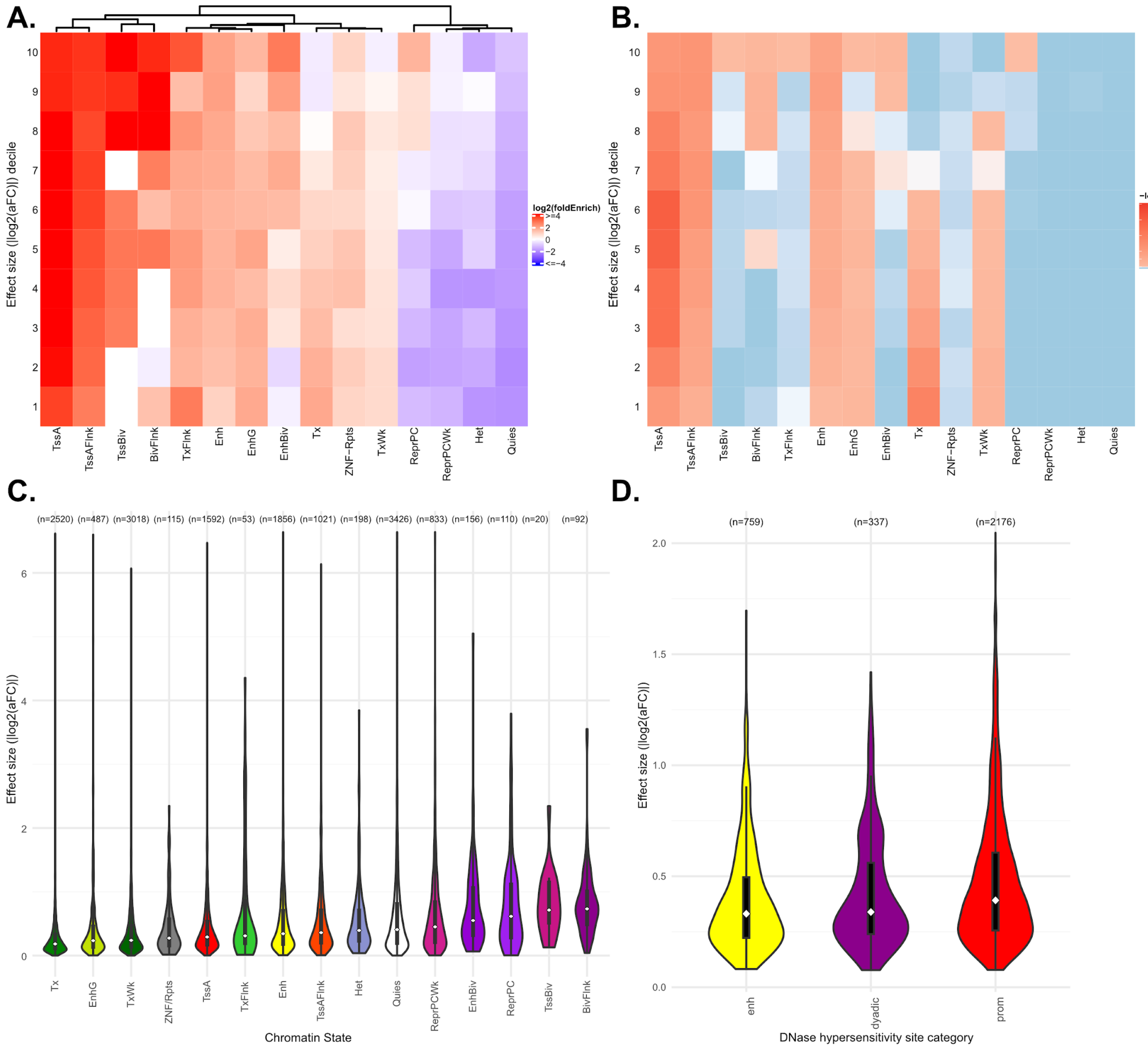


**Figure S13. Lead eQTLs are functionally enriched in the regulatory regions of primary B-cells (E032). (A)** Decile-Based enrichment analysis of lead eQTL effect sizes measured as absolute value of the estimated allelic fold change ($|log_2(aFC)|$) across 15 different chromatin states predicted by the Roadmap Epigenomics chromHMM model[30] specific to Primary B-Cells. Heatmap showcases promoter associated regions maintaining a consistent trend across all deciles of eQTL effect sizes. In contrast, a notable enrichment is detected within bivalent regions, Bivalent Transcription Start Sites (TSSBiv), Bivalent Enhancers (EnhBiv) and Bivalent flanking regions (BivFlnk), which is most pronounced for eQTLs demonstrating large effects. **(B)** Corresponding heatmap to panel A, showing significance of the enrichment estimates (right tailed, Binomial P-value). **(C)** Distribution of effect sizes for lead eQTLs within chromatin states exhibiting pronounced trend of diminished effect sizes for transcriptional elongation regions (Tx, TxWk, and TxFlnk) **(D)** Distribution of absolute effect sizes of lead eQTLs (measured as $log_2(aFC)$) across chromatin states in Primary B-Cells that are associated with multi-tissue DNAse hypersensitivity sites[30].

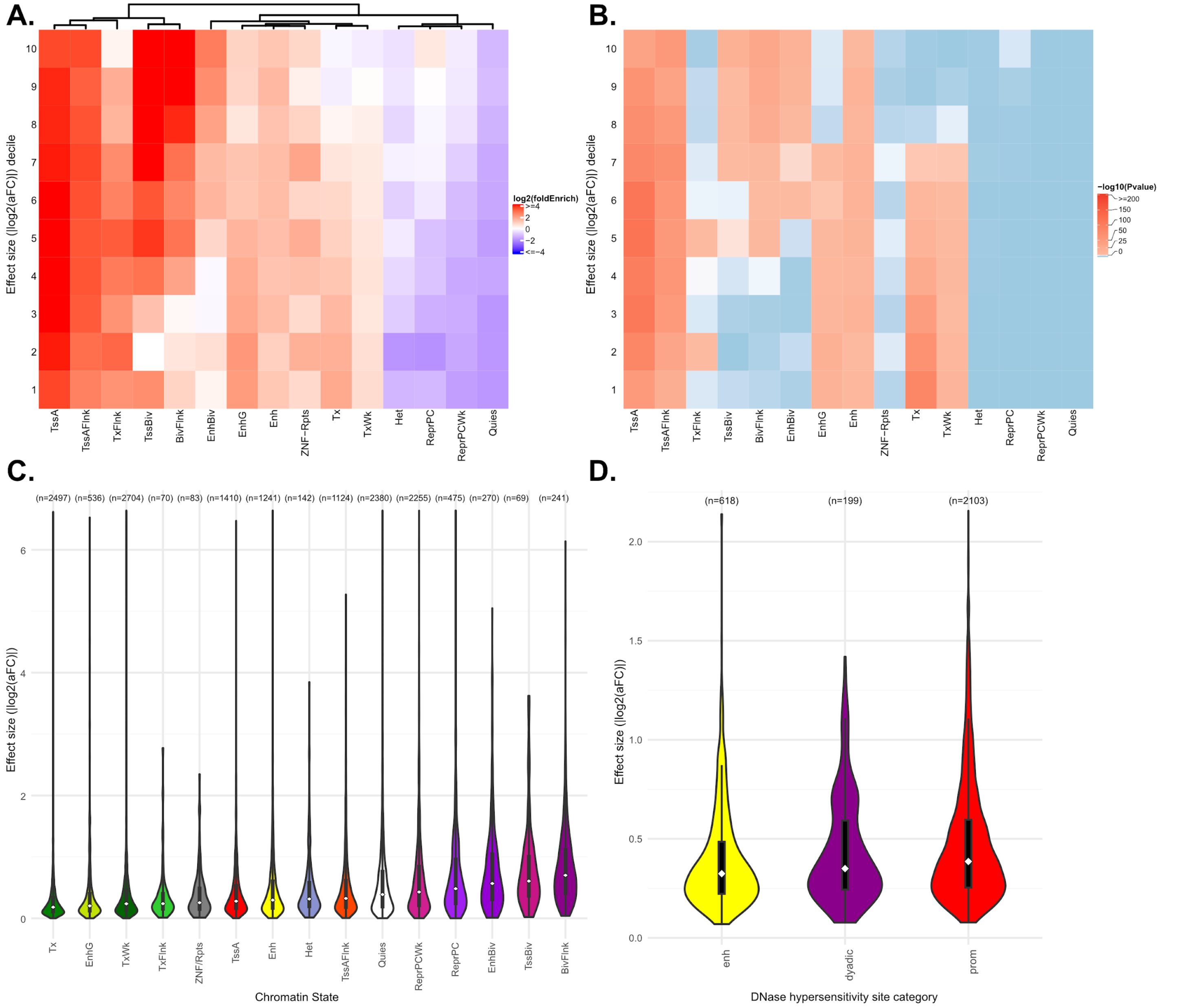


**Figure S14. Lead eQTLs are functionally enriched in the regulatory regions of primary T-cells (E034). (A)** Decile-Based enrichment analysis of lead eQTL effect sizes measured as absolute value of the estimated allelic fold change ($|\log_2(\text{aFC})|$) across 15 different chromatin states predicted by chromHMM model specific to Primary T-Cells . Heatmap showcases promoter associated regions maintaining a consistent trend across all deciles of eQTL effect sizes. In contrast, a notable enrichment is detected within bivalent regions, Bivalent Transcription Start Sites (TSSBiv), Bivalent Enhancers (EnhBiv) and Bivalent flanking regions (BivFlnk), which is most pronounced for eQTLs demonstrating large effects. **(B)** Corresponding heatmap to panel A, showing significance of the enrichment estimates (right tailed, Binomial P-value). **(C)** Distribution of effect sizes for lead eQTLs within chromatin states exhibiting pronounced trend of diminished effect sizes for transcriptional elongation regions (Tx, TxWk, and TxFlnk) **(D)** Distribution of absolute effect sizes of lead eQTLs measured as $\log_2(\text{aFC})$ across chromatin states in Primary T-Cells that are associated with multi-tissue DNAse Hypersensitivity Sites[30].

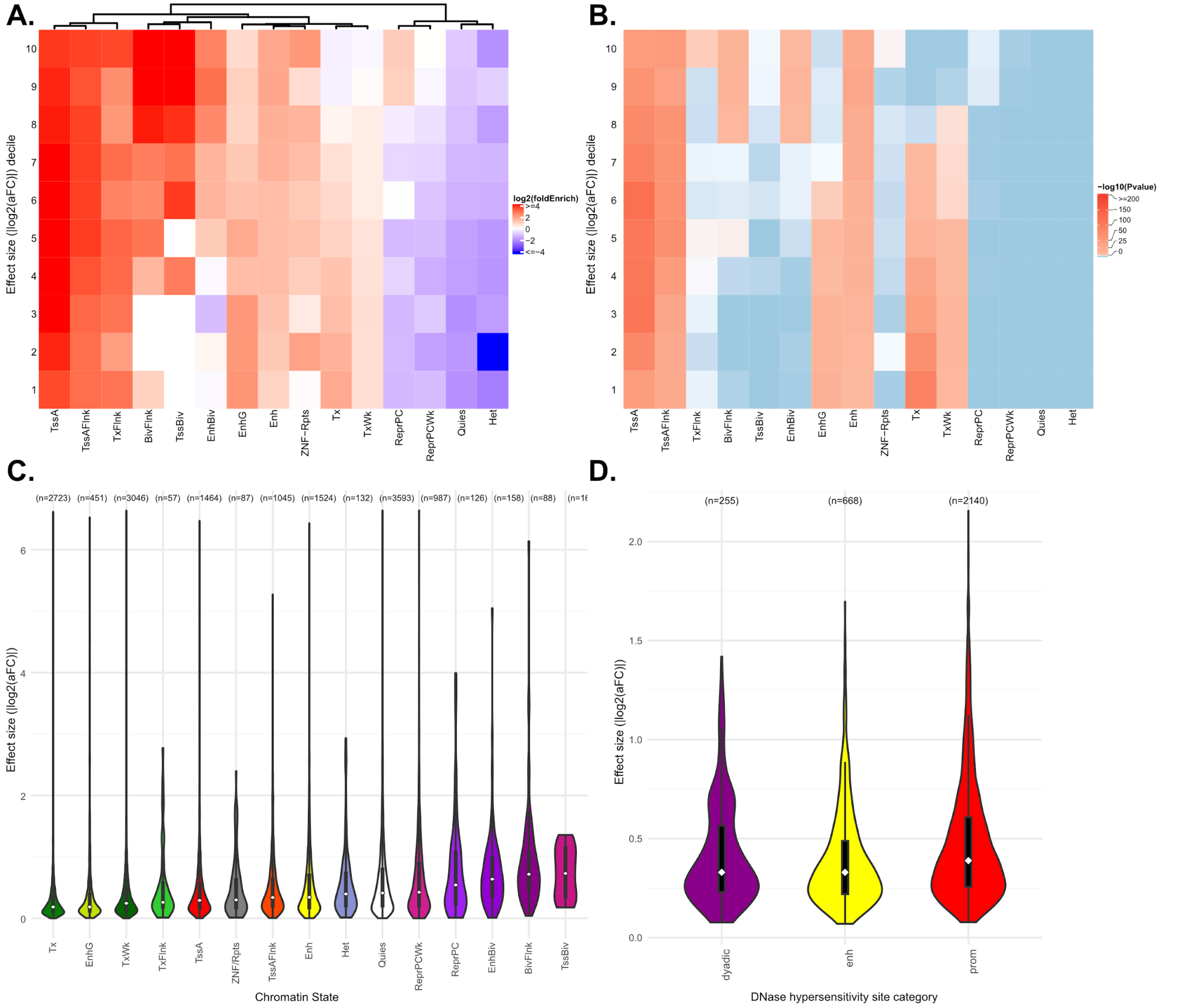


**Figure S15. Lead eQTLs are functionally enriched in the regulatory regions of primary natural killer cells (E046). (A)** Decile-Based enrichment analysis of lead eQTL effect sizes measured as absolute value of the estimated allelic fold change ($|\log_2(\text{aFC})|$) across 15 different chromatin states predicted by chromHMM model specific to Primary Natural Killer Cells. Heatmap showcases promoter associated regions maintaining a consistent trend across all deciles of eQTL effect sizes. In contrast, a notable enrichment is detected within bivalent regions, Bivalent Enhancers (EnhBiv) and Bivalent flanking regions (BivFlnk), which is most pronounced for eQTLs demonstrating large effects. **(B)** Corresponding heatmap to panel A, showing significance of the enrichment estimates (right tailed, Binomial P-value). **(C)** Distribution of effect sizes for lead eQTLs within chromatin states exhibiting pronounced trend of diminished effect sizes for transcriptional elongation regions (Tx, TxWk, and TxFlnk) **(D)** Distribution of absolute effect sizes of lead eQTLs measured as $\log_2(\text{aFC})$ across chromatin states in Primary Natural Killer Cells that are associated with multi-tissue DNAse Hypersensitivity Sites[30].

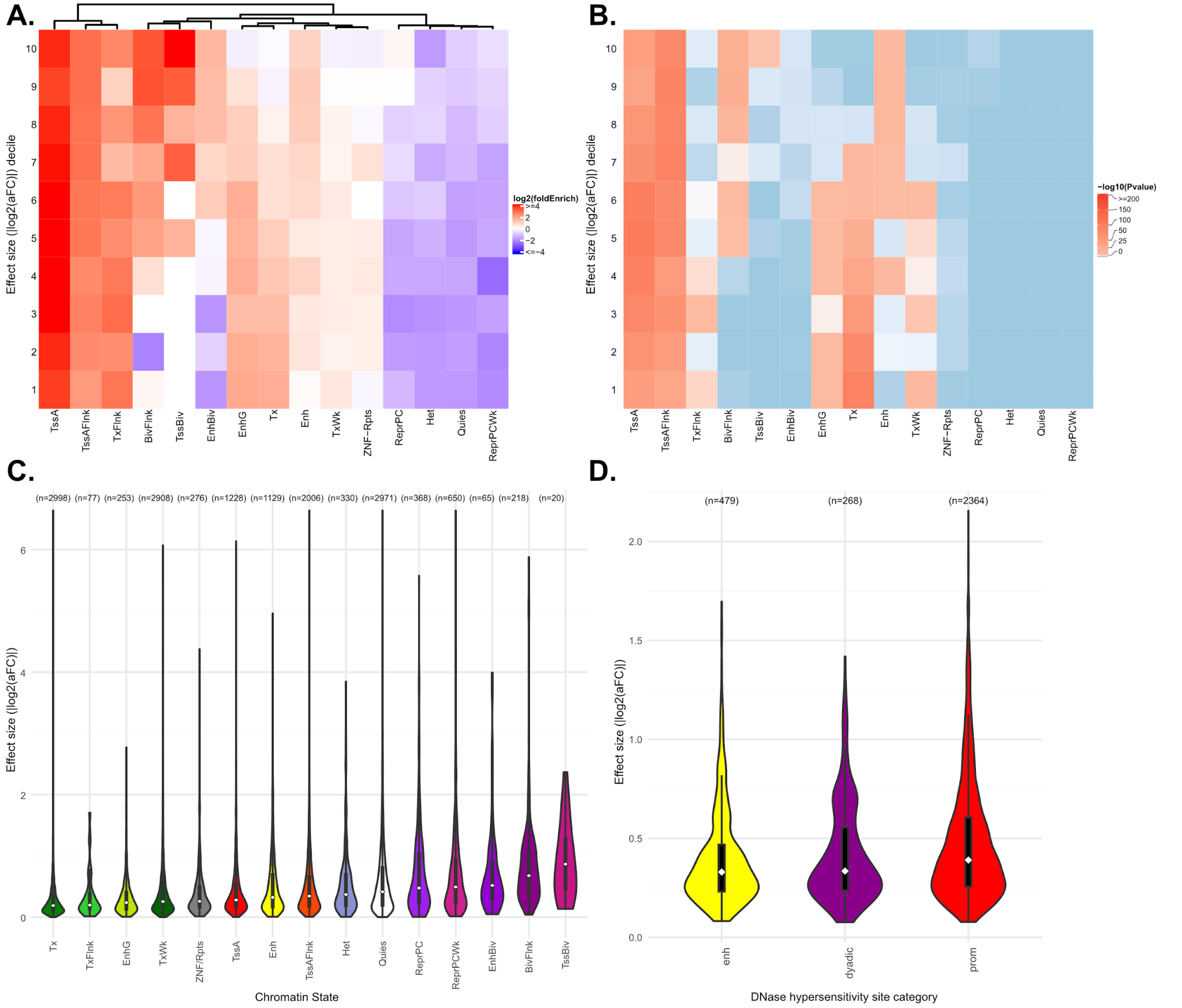


**Figure S16. Lead eQTLs are functionally enriched in the regulatory regions of primary hematopoietic stem cells (E051). (A)** Decile-Based enrichment analysis of lead eQTL effect sizes measured as absolute value of the estimated allelic fold change ($|\log_2(aFC)|$) across 15 different chromatin states predicted by chromHMM model specific to Primary Hematopoietic Stem Cells. Heatmap showcases promoter associated regions maintaining a consistent trend across all deciles of eQTL effect sizes. In contrast, a notable enrichment is detected within bivalent regions, Bivalent Transcription Start Sites (TSSBiv) and Bivalent flanking regions (BivFlnk), which is most pronounced for eQTLs demonstrating large effects. **(B)** Corresponding heatmap to panel A, showing significance of the enrichment estimates (right tailed, Binomial P-value). **(C)** Distribution of effect sizes for lead eQTLs within chromatin states exhibiting pronounced trend of diminished effect sizes for transcriptional elongation regions (Tx, TxWk, and TxFlnk) **(D)** Distribution of absolute effect sizes of lead eQTLs measured as $\log_2(aFC)$ across chromatin states in Primary Hematopoietic Stem Cells that are associated with multi-tissue DNAse Hypersensitivity Sites[30].

The enrichment calculations for both chromatin states and DHS peaks were conducted using the same GREGOR Perl script pipeline[72], as previously applied in the transcription factor binding site enrichment analysis. The enrichment was quantified using $\log_2$ fold changes (observed/expected) and p-values ($-\log_{10}$ transformed) to determine the magnitude and significance of enrichment across chromatin states and DHS sites across 127 cell/tissue samples.

### 14 Lead e- and sQTL AF differentiation between populations

To visualize the joint frequency spectrum of lead eQTLs, we used the GeoVar software package[74]. For visualization, we defined the following discrete allele frequency categories for visualization based on the within-dataset alternative allele frequency: unobserved (allele frequency = 0%), rare (0% < allele frequency < 5%), and common (allele frequency > 5%). All allele frequencies were calculated using bcftools (version 1.17). We used GeoVar to visualize the joint frequency spectrum of all lead eQTLs (**Fig. S17A**), lead eQTLs that are rare or unobserved in at least one continental groups (**Fig. S17B**), lead eQTLs that are unobserved in the EUR continental group (**Fig. S17C**), and lead eQTLs that are unobserved in both EUR and AFR continental groups (**Fig. S17D**).

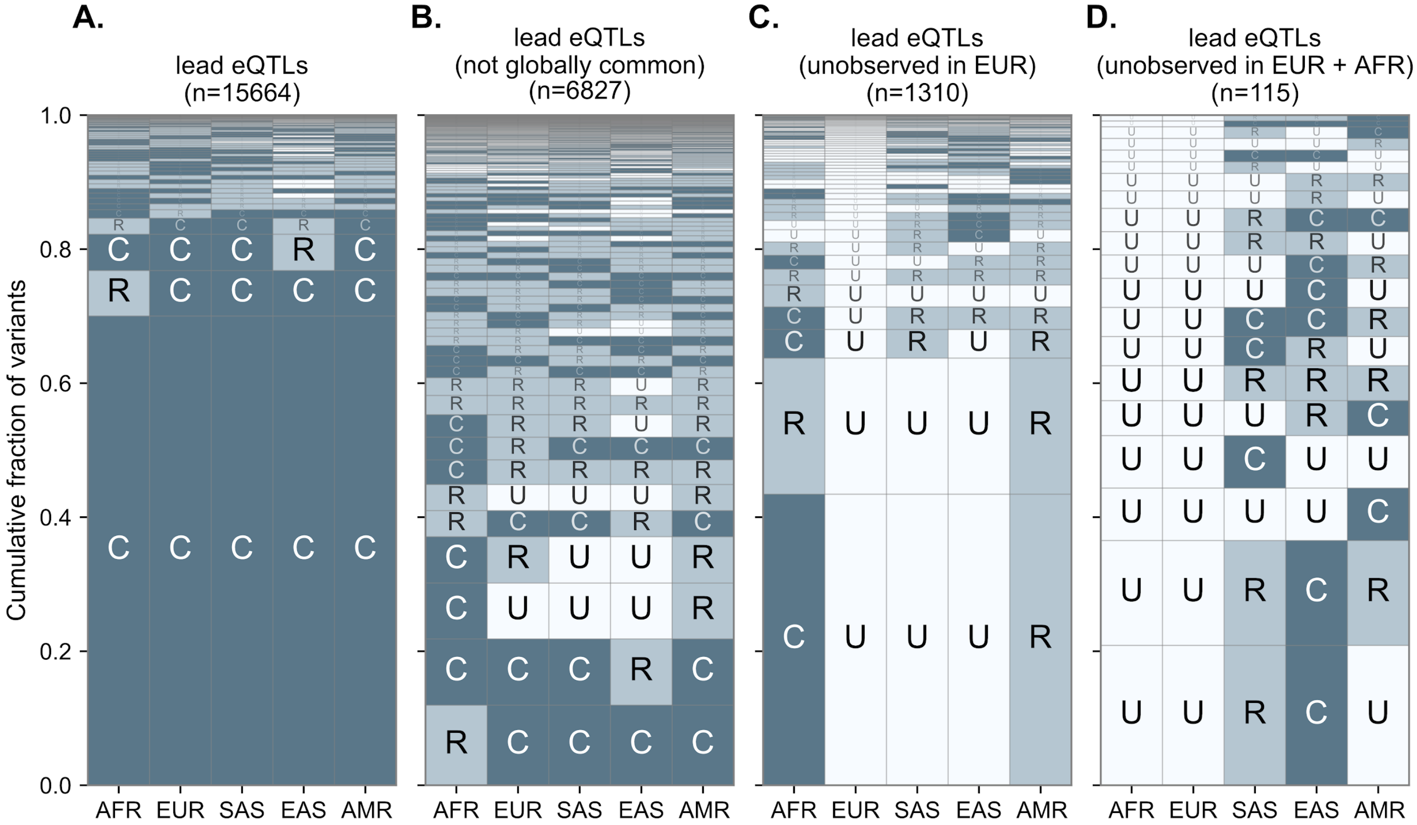


**Figure S17. Population stratification of eQTLs.** Geographic frequencies of lead eQTLs found in MAGE across **(A)** all lead eQTLs, **(B)** excluding variants with allele frequencies > 5% across all regional populations, **(C)** only including variants unobserved in European ancestry populations, and **(D)** only including variants unobserved in both European and African-ancestry populations. The geographic distributions are sorted by the most common at the bottom and rarest at the top. Allele frequencies are categorized as unobserved (U), rare variants with population allele frequencies < 5% (R), and common variants (C) with allele frequencies greater than 5%.

We also investigated the joint frequency spectrum of gene-level lead sQTLs (**Fig. S18**).

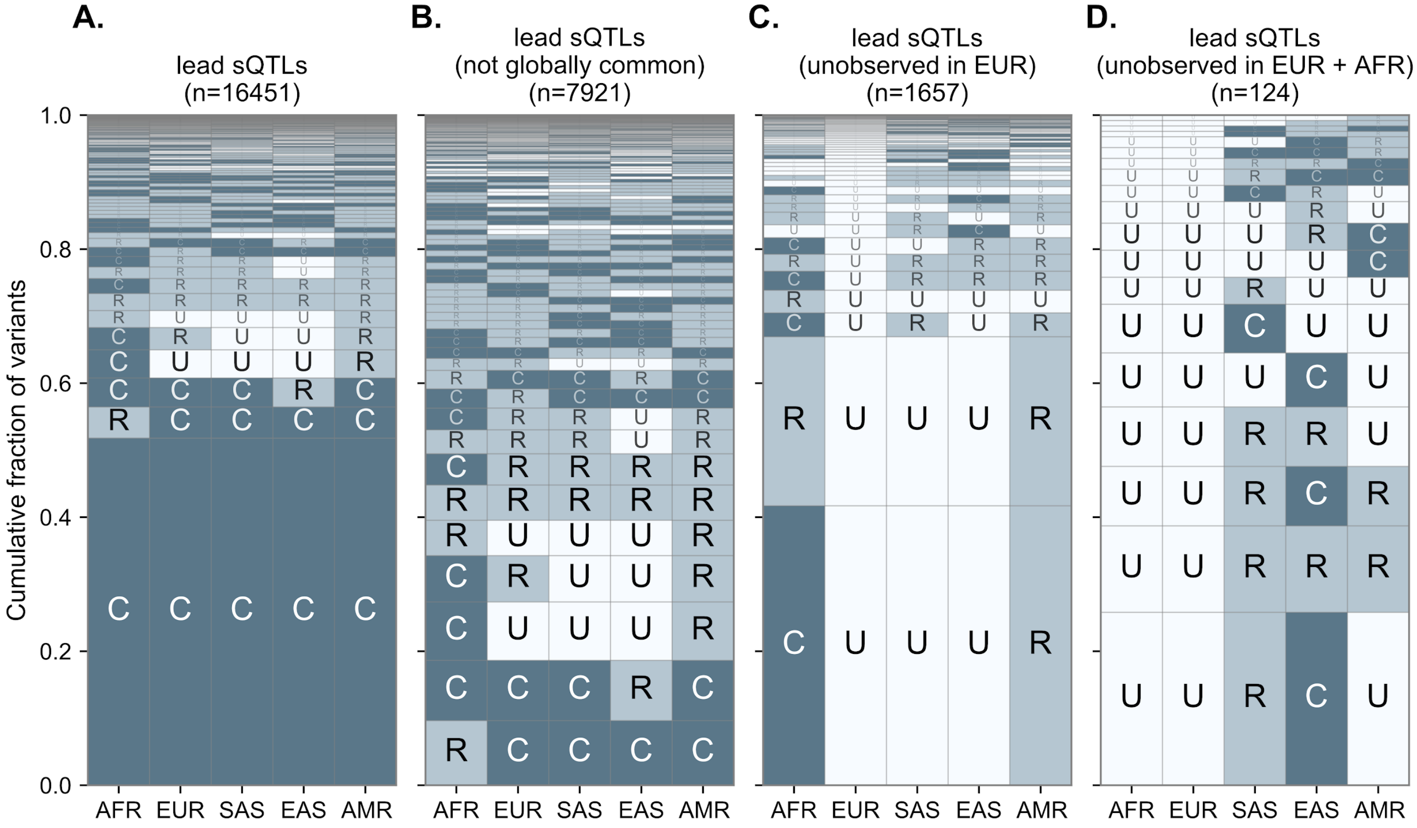


**Figure S18. Population stratification of sQTLs.** Geographic frequencies of lead sQTLs found in MAGE across **(A)** all lead sQTLs, **(B)** excluding variants with allele frequency > 5% (i.e., "globally common") across all regional populations, **(C)** only including variants unobserved in European ancestry populations, and **(D)** only including variants unobserved in both European and African-ancestry populations. Allele frequencies are categorized as unobserved (U), rare variants with population allele frequencies < 5% (R), and common variants (C) with allele frequencies greater than 5%.

### 15 Replication of credible sets in GTEx

#### 15.1 Defining replicating vs. non-replicating eQTLs

To appropriately compare *cis*-eQTL signals between MAGE and GTEx[15], we first collapsed tissue-specific DAP-G credible sets from GTEx into cross-tissue merged credible sets (ctmCS) for each gene. To construct the ctmCS, for each gene, we combined the fine-mapping credible sets inferred using DAP-G across all tissues for that specific gene (restricting to variants with PIP > 0.95)[75]. To combine the per-tissue credible sets we iteratively joined any DAP-G credible sets sharing variants, resulting in a set of non-overlapping variants per ctmCS per gene. We considered a MAGE credible set to replicate in GTEx if any variant contained in the MAGE credible set was also contained in at least one GTEx ctmCS for that gene. To define the lead eQTL within each GTEx ctmCS, we first select the tissue-level credible set with the highest coverage amongst those that comprise the ctmCS, and from that tissue-level credible set, we select the variant with the highest PIP. We observe that the set of GTEx ctmCS's that do not replicate in MAGE is enriched for ctmCS's that comprise tissue-level CS's from only a single tissue (hence are tissue-specific; **Fig. S19**).

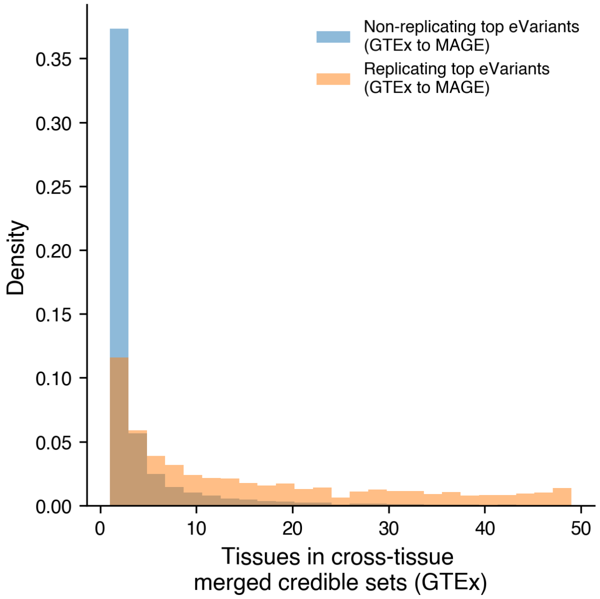


**Figure S19. GTEx DAP-G fine-mapping signals that do not replicate in MAGE are largely tissue-specific.** Comparison of number of tissues contained by 79,913 cross-tissue merged credible sets (ctmCS) from GTEx that do not replicate in MAGE against 7,913 ctmCS that replicate in MAGE. The number of tissues is defined as the number of tissues across all variants included in a ctmCS.

#### 15.2 Functional annotation and enrichment of non-replicating eQTLs

Because so many of the MAGE lead eQTLs did not replicate in the GTEx fine-mapping results, we were acutely curious whether this subset of our results was enriched for functional variation. To address this question, we repeated the analysis described in section **13** for the subset of MAGE eQTLs that did not replicate in the GTEx DAP-G results. Briefly, to evaluate chromatin context of MAGE specific eQTLs, we performed the enrichment and functional annotation analysis across all the 15 predicted chromatin states in 127 epigenomic mappings and 53 primary DHS data from Roadmap Epigenomics as described insection **13**. Our findings mirrored previous results, demonstrating similar functional enrichment patterns between the non-replicating subset of MAGE eQTLs and the full set of results (**Fig. S20A-C**). Additionally, to examine the relationship between fine-mapped MAGE-specific lead eQTLs and the binding sites of transcription factors (TFs) and chromatin regulators (CRs), we conducted a detailed enrichment analysis with 338 DNA associated ChIP-seq profiles obtained from the ENCODE Project Consortium as described in section **13**. As with the full set of lead eQTLs, we observed that MAGE-specific lead eQTLs were highly enriched in TF binding site annotations (**Fig. S20D**).

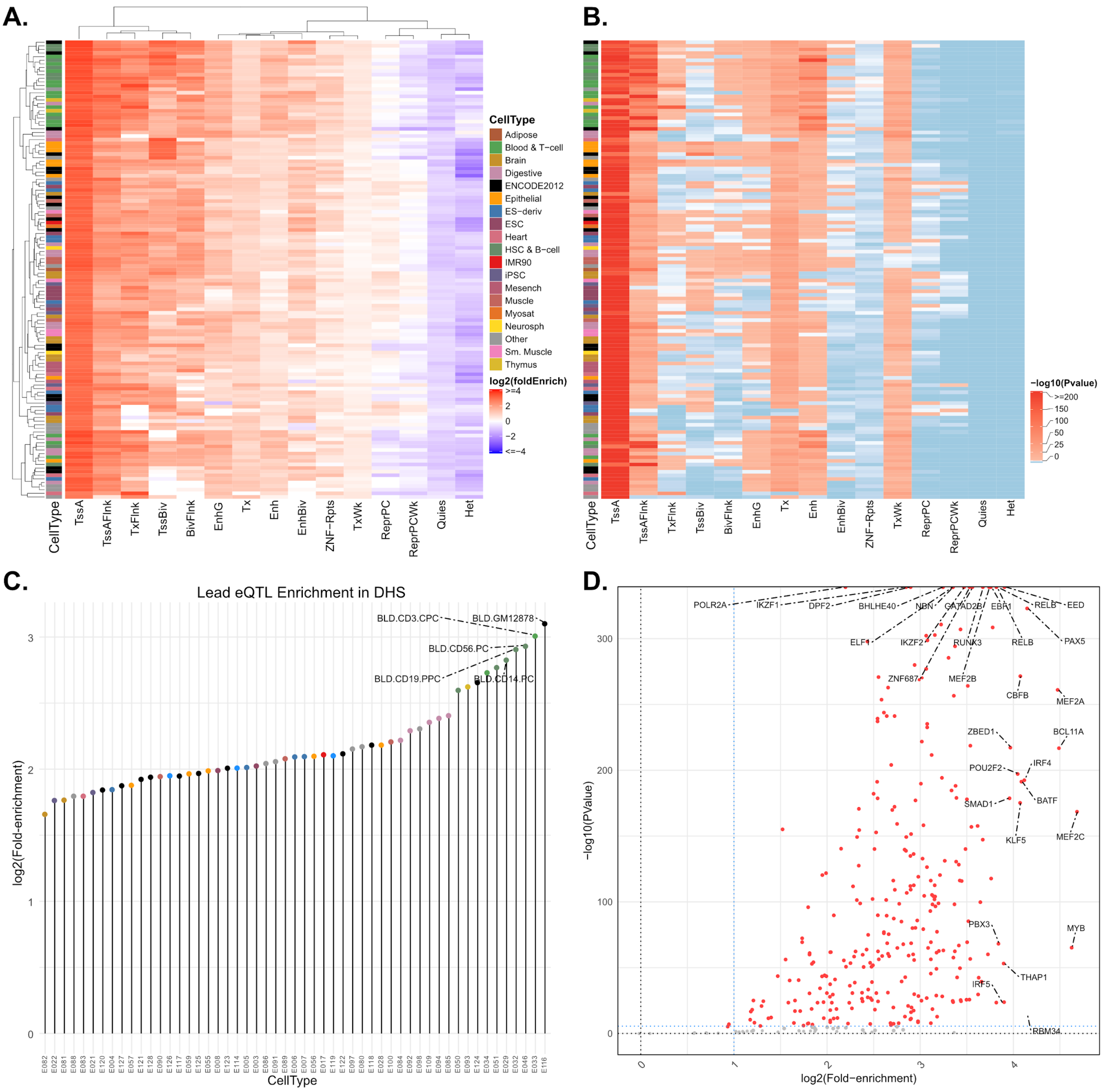

**Figure S20. Lead eQTLs that did not replicate in GTEx are functionally enriched in the regulatory regions of pertinent cell types. (A)** A heatmap showing the enrichment of unique lead eQTLs at 15 predicted chromatin states across 127 cell/tissue types from the Roadmap Epigenomic Consortium. Cell type annotations are displayed by colored legend keys. Lead eQTLs exhibited strong enrichment at promoter (TssA, TssAFlnk) and enhancer regions (Enh, EnhG) both, with promoter regions showing more pronounced enrichment compared to enhancer regions across the cell types. Strong Hierarchical clustering of Blood and T cells highlighted by green colored cell type annotation bar on the left. **(B)** Corresponding heatmap to panel A, showing significance of the enrichment estimates (right-tailed, Binomial P-value). **(C)** A lollipop plot showing the pronounced enrichment of lead eQTLs in the DHSs (DNAse Hypersensitivity Sites) across 53 cell/tissue types (colored as in A). We note a marked enrichment in DHS of blood cell types with lymphoblastoid cell line GM12878 as one of the top hits. **(D)** Volcano plot representing the enrichment analysis of lead eQTLs at TFBSs (Transcription factor binding sites) of 338 TF ChIP-seq binding profiles sourced from ENCODE. Data points reflecting a

Bonferroni-corrected p-value < 0.001 and log2(fold-enrichment) > 1 stand out in red, underscoring those transcription factors where lead cis-eQTL enrichment is both statistically significant and of notable magnitude.

Functional annotation of one such eQTL signal that did not replicate in GTEx is shown in **Fig. S21**. The variant rs115070172 is associated with decreased expression of *GSTP1* and is largely private to the AMR continental group, as shown in **Fig. 4B**[76].

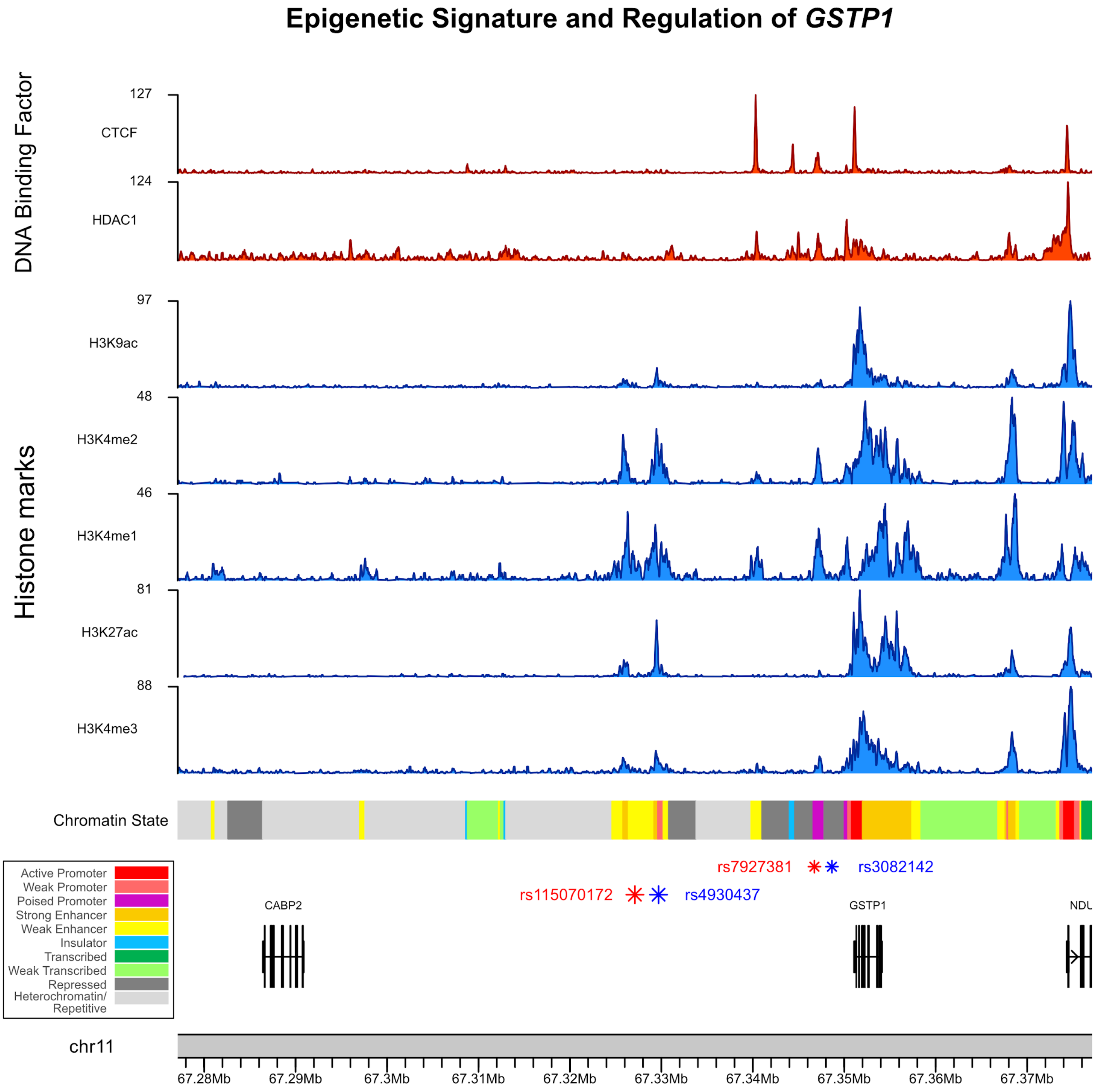

**Figure S21. Functional epigenetic annotation of *GSTP1* eQTL credible signals.** ENCODE epigenetic signals at the *GSTP1* locus, within lymphoblastoid cell line GM12878. From top down: 1) Red colored tracks denote binding of DNA associated factors. 2) Blue colored tracks show histone mark signals, including promoter and enhancer associated chromatin marks. 3) The multi-colored track shows the predicted chromatin state along the chromosome, legend at bottom left. 4) Asterisks represent fine-mapped eQTLs for *GSTP1*. The two red asterisks represent the lead eQTLs of the two *GSTP1* credible sets and the blue asterisks represent the corresponding SNP with the next highest PIP within the same credible set. The top two (smaller) asterisks represent eQTLs from one credible set, the bottom two (larger) asterisks represent eQTLs from the other credible set (which was highlighted in **Fig. 5**). 5) Gene annotation at and around the *GSTP1* locus.

### 16 Relationship between fixation index and differential gene expression

Weir & Cockerham's $F_{ST}$[39] was calculated for each fine-mapped lead eQTL identified in section **10.5** using the statistic's implementation in vcflib (version 1.0.0_rc2)[77]. For each lead eQTL, $F_{ST}$ was estimated for each of the five target continental groups, where foreground samples fell within the target population (e.g., AFR) and background samples fell within any of the remaining four continental groups (e.g., EUR, SAS, EAS, or AMR). An average $F_{ST}$ per eGene was calculated for each population by computing the mean of all eQTL $F_{ST}$'s identified for each respective eGene (negative $F_{ST}$ values were converted to zeroes).

### 17 *cis*-eQTL effect size heterogeneity between populations

We discovered *cis*-eQTLs exhibiting effect size heterogeneity across continental groups (he-eQTLs) from among the lead eQTLs from each SuSiE credible set (described in section **10.5**). To ensure that we are detecting robust signals, we first filter to only those variants with MAF ≥ 0.05 in at least two continental groups. After filtering, 12,338/15,664 credible sets remain for analysis. From this set, we used two similar yet distinct approaches to discover he-eQTLs.

In the first approach, for each lead eQTL we fit two models. We first fit a model regressing normalized TMM values onto sample genotype and eQTL mapping covariates (sex, top 5 genotyping PCs, 60 PEER factors). This is described in model **[1]** below, and we note that this is equivalent to the model used for nominal eQTL mapping with FastQTL (described in section **10.4**). Here, $g_{j,i}$ describes the sample genotypes at the top variant of the $i^{th}$ credible set for gene $j$, $E_j$ describes the inverse normal transformed TMM values of gene $j$, and $X_{sex}$, $X_{PCA}$, and $X_{PEER}$ describe the covariates used for mapping. Next, we fit a model identical to model **[1]** but that now includes an additional genotype-by-continental group interaction term, as described in model **[2]**. Here, $X_{CG}$ describes the continental groups of the samples. We performed an F-test to determine if the more complex model **[2]** explains the data significantly better than model **[1]**. We define he-eQTLs as those variants with significant F-statistics after Bonferroni correction ($p < 4 \times 10^{-6}$).

$$\textbf{(1)}\quad E_j \sim g_{j,i} + X_{sex} + X_{PCA} + X_{PEER}$$

$$\textbf{(2)}\quad E_j \sim g_{j,i} + (g_{j,i} \times X_{CG}) + X_{sex} + X_{PCA} + X_{PEER}$$

The second approach mirrors the first with one important distinction: for each gene with multiple causal signals (i.e. multiple SuSiE credible sets), all top hit variants for that gene were included in the regression. This effectively controls for the additive effects of multiple causal SNPs. So, for each SuSiE top hit variant, we first fit a model regressing normalized TMM values onto sample genotypes for the focal top hit variant and all other top hit variants for that gene (regardless of MAF), along with eQTL mapping covariates. This is described in model **[3]** below, assuming $n$ credible sets for the focal gene. All variables are as described above, with $g_{j,i}$ being the genotypes of the focal variant. We next fit a model identical to model **[3]** but that now includes an additional genotype-by-continental group interaction term *for the focal variant only*. This is described in model **[4]**. As before, we performed an F-test to determine if model **[4]** explains the data significantly better than model **[3]**, and we define he-eQTLs as those variants with significant F-statistics after Bonferroni correction ($p$ < 4E-6).

$$\textbf{(3)}\ \ E_j \sim g_{j,1} + \ldots + g_{j,i} + \ldots + g_{j,n} + X_{sex} + X_{PCA} + X_{PEER}$$

$$\textbf{(4)}\ \ E_j \sim g_{j,1} + \ldots + g_{j,i} + \ldots + g_{j,n} + (g_{j,i} \times X_{CG}) + X_{sex} + X_{PCA} + X_{PEER}$$

Models 1-4 were fit using the `formula.api.ols` function from the *statsmodels* package (version 0.14.0) in Python. The F-test between models 1 and 2 and between models 3 and 4 was performed using the `stats.anova.anova_lm` function from the *statsmodels* package.

### References:


1. Li, Y. I. *et al.* RNA splicing is a primary link between genetic variation and disease. *Science* **352**, 600–604 (2016).
2. Brem, R. B., Yvert, G., Clinton, R. & Kruglyak, L. Genetic dissection of transcriptional regulation in budding yeast. *Science* **296**, 752–755 (2002).
3. Morley, M. *et al.* Genetic analysis of genome-wide variation in human gene expression. *Nature* **430**, 743–747 (2004).
4. GTEx Consortium. Genetic effects on gene expression across human tissues. *Nature* **550**, 204–213 (2017).
5. Mogil, L. S. *et al.* Genetic architecture of gene expression traits across diverse populations. *PLoS Genet.* **14**, e1007586 (2018).
6. Lappalainen, T. *et al.* Transcriptome and genome sequencing uncovers functional variation in humans. *Nature* **501**, 506–511 (2013).
7. Popejoy, A. B. & Fullerton, S. M. Genomics is failing on diversity. *Nature* **538**, 161–164 (2016).
8. Martin, A. R. *et al.* Human Demographic History Impacts Genetic Risk Prediction across Diverse Populations. *Am. J. Hum. Genet.* **107**, 788–789 (2020).
9. Wojcik, G. L. *et al.* Genetic analyses of diverse populations improves discovery for complex traits. *Nature* **570**, 514–518 (2019).
10. Kita, R., Venkataram, S., Zhou, Y. & Fraser, H. B. High-resolution mapping of cis-regulatory variation in budding yeast. *Proc Natl Acad Sci U S A* . **114**, E10736–E10744 (2017).
11. 1000 Genomes Project Consortium *et al.* A global reference for human genetic variation. *Nature* **526**, 68–74 (2015).
12. DeGorter, M. K. *et al.* Transcriptomics and chromatin accessibility in multiple African population samples. *bioRxiv*.
13. International HapMap Consortium. The International HapMap Project. *Nature* **426**, 789–796 (2003).
14. Alexander, D. H., Novembre, J. & Lange, K. Fast model-based estimation of ancestry in unrelated individuals. *Genome Res.* **19**, 1655–1664 (2009).
15. GTEx Consortium. The GTEx Consortium atlas of genetic regulatory effects across human tissues. *Science* **369**, 1318–1330 (2020).
16. Li, Y. I. *et al.* Annotation-free quantification of RNA splicing using LeafCutter. *Nat. Genet.* **50**, 151–158 (2018).

17. Lewontin, R. C. The apportionment of human diversity. in *Evolutionary Biology* 381–398 (Springer US, 1972).

18. Jorde, L. B. *et al.* The distribution of human genetic diversity: a comparison of mitochondrial, autosomal, and Y-chromosome data. *Am. J. Hum. Genet.* **66**, 979–988 (2000).

19. Martin, A. R. *et al.* Transcriptome sequencing from diverse human populations reveals differentiated regulatory architecture. *PLoS Genet.* **10**, e1004549 (2014).

20. Bergström, A. *et al.* Insights into human genetic variation and population history from 929 diverse genomes. *Science* **367**, (2020).

21. Ramachandran, S. *et al.* Support from the relationship of genetic and geographic distance in human populations for a serial founder effect originating in Africa. *Proc. Natl. Acad. Sci. U. S. A.* **102**, 15942–15947 (2005).

22. Prugnolle, F., Manica, A. & Balloux, F. Geography predicts neutral genetic diversity of human populations. *Curr. Biol.* **15**, R159–60 (2005).

23. Byrska-Bishop, M. *et al.* High-coverage whole-genome sequencing of the expanded 1000 Genomes Project cohort including 602 trios. *Cell* **185**, 3426–3440.e19 (2022).

24. Zou, Y., Carbonetto, P., Wang, G. & Stephens, M. Fine-mapping from summary data with the 'Sum of Single Effects' model. *PLoS Genet.* **18**, e1010299 (2022).

25. Jansen, R. *et al.* Conditional eQTL analysis reveals allelic heterogeneity of gene expression. *Hum. Mol. Genet.* **26**, 1444–1451 (2017).

26. Mohammadi, P., Castel, S. E., Brown, A. A. & Lappalainen, T. Quantifying the regulatory effect size of cis-acting genetic variation using allelic fold change. *Genome Res.* **27**, 1872–1884 (2017).

27. Huang, Q. Q., Ritchie, S. C., Brozynska, M. & Inouye, M. Power, false discovery rate and Winner's Curse in eQTL studies. *Nucleic Acids Res.* **46**, e133 (2018).

28. Lek, M. *et al.* Analysis of protein-coding genetic variation in 60,706 humans. *Nature* **536**, 285–291 (2016).

29. Glassberg, E. C., Gao, Z., Harpak, A., Lan, X. & Pritchard, J. K. Evidence for Weak Selective Constraint on Human Gene Expression. *Genetics* **211**, 757–772 (2019).

30. Roadmap Epigenomics Consortium *et al.* Integrative analysis of 111 reference human epigenomes. *Nature* **518**, 317–330 (2015).

31. ENCODE Project Consortium. An integrated encyclopedia of DNA elements in the human genome. *Nature* **489**, 57–

74 (2012).

32. Patel, R. A. *et al.* Genetic interactions drive heterogeneity in causal variant effect sizes for gene expression and complex traits. *Am. J. Hum. Genet.* **109**, 1286–1297 (2022).

33. Hou, K. *et al.* Causal effects on complex traits are similar for common variants across segments of different continental ancestries within admixed individuals. *Nat. Genet.* **55**, 549–558 (2023).

34. Visscher, P. M. *et al.* 10 Years of GWAS Discovery: Biology, Function, and Translation. *Am. J. Hum. Genet.* **101**, 5–22 (2017).

35. Gutenkunst, R. N., Hernandez, R. D., Williamson, S. H. & Bustamante, C. D. Inferring the Joint Demographic History of Multiple Populations from Multidimensional SNP Frequency Data. *PLoS Genet.* **5**, (2009).

36. Fang, C. *et al.* Aberrant GSTP1 promoter methylation is associated with increased risk and advanced stage of breast cancer: a meta-analysis of 19 case-control studies. *BMC Cancer* **15**, 1–8 (2015).

37. Louie, S. M. *et al.* GSTP1 Is a Driver of Triple-Negative Breast Cancer Cell Metabolism and Pathogenicity. *Cell Chemical Biology* **23**, 567–578 (2016).

38. Arai, T. *et al.* Association of GSTP1 CpG Islands Hypermethylation with Poor Prognosis in Human Breast Cancers. *Breast Cancer Res. Treat.* **100**, 169–176 (2006).

39. Weir, B. S. & Cockerham, C. C. ESTIMATING F-STATISTICS FOR THE ANALYSIS OF POPULATION STRUCTURE. *Evolution* **38**, 1358–1370 (1984).

40. Saitou, M., Dahl, A., Wang, Q. & Liu, X. Allele frequency differences of causal variants have a major impact on low cross-ancestry portability of PRS. *medRxiv* 2022.10.21.22281371 (2022) doi:10.1101/2022.10.21.22281371.

41. Kachuri, L. *et al.* Gene expression in African Americans, Puerto Ricans and Mexican Americans reveals ancestry-specific patterns of genetic architecture. *Nat. Genet.* **55**, 952–963 (2023).

42. Mostafavi, H., Spence, J. P., Naqvi, S. & Pritchard, J. K. Systematic differences in discovery of genetic effects on gene expression and complex traits. *Nat. Genet.* (2023) doi:10.1038/s41588-023-01529-1.

43. Rau, C. D. *et al.* Modeling epistasis in mice and yeast using the proportion of two or more distinct genetic backgrounds: Evidence for ‘polygenic epistasis’. *PLoS Genet.* **16**, e1009165 (2020).

44. Cheung, V. G. *et al.* Natural variation in human gene expression assessed in lymphoblastoid cells. *Nat. Genet.* **33**, 422–425 (2003).

45. Strober, B. J. *et al.* Dynamic genetic regulation of gene expression during cellular differentiation. *Science* **364**, 1287–1290 (2019).

46. Workman, R. E. *et al.* Nanopore native RNA sequencing of a human poly(A) transcriptome. *Nat. Methods* **16**, 1297–1305 (2019).

47. Glinos, D. A. *et al.* Transcriptome variation in human tissues revealed by long-read sequencing. *Nature* **608**, 353–359 (2022).

48. Reese, F. *et al.* The ENCODE4 long-read RNA-seq collection reveals distinct classes of transcript structure diversity. *bioRxiv* (2023) doi:10.1101/2023.05.15.540865.

49. Sibbesen, J. A. *et al.* Haplotype-aware pantranscriptome analyses using spliced pangenome graphs. *Nat. Methods* **20**, 239–247 (2023).

50. Claw, K. G. *et al.* A framework for enhancing ethical genomic research with Indigenous communities. *Nat. Commun.* **9**, 2957 (2018).

51. Purcell, S. *et al.* PLINK: a tool set for whole-genome association and population-based linkage analyses. *Am. J. Hum. Genet.* **81**, 559–575 (2007).

52. Frankish, A. *et al.* GENCODE reference annotation for the human and mouse genomes. *Nucleic Acids Res.* **47**, D766–D773 (2019).

53. Patro, R., Duggal, G., Love, M. I., Irizarry, R. A. & Kingsford, C. Salmon provides fast and bias-aware quantification of transcript expression. *Nat. Methods* **14**, 417–419 (2017).

54. Soneson, C., Love, M. I. & Robinson, M. D. Differential analyses for RNA-seq: transcript-level estimates improve gene-level inferences. *F1000Res.* **4**, 1521 (2015).

55. Dobin, A. *et al.* STAR: ultrafast universal RNA-seq aligner. *Bioinformatics* **29**, 15–21 (2013).

56. Fairley, S., Lowy-Gallego, E., Perry, E. & Flicek, P. The International Genome Sample Resource (IGSR) collection of open human genomic variation resources. *Nucleic Acids Res.* **48**, D941–D947 (2019).

57. Garrido-Martín, D., Calvo, M., Reverter, F. & Guigó, R. A fast non-parametric test of association for multiple traits. *bioRxiv* 2022.06.06.493041 (2022) doi:10.1101/2022.06.06.493041.

58. Love, M. I., Huber, W. & Anders, S. Moderated estimation of fold change and dispersion for RNA-seq data with DESeq2. *Genome Biol.* **15**, 550 (2014).

59. Robinson, M. D. & Oshlack, A. A scaling normalization method for differential expression analysis of RNA-seq data. *Genome Biol.* **11**, R25 (2010).

60. Robinson, M. D., McCarthy, D. J. & Smyth, G. K. edgeR: a Bioconductor package for differential expression analysis of digital gene expression data. *Bioinformatics* **26**, 139–140 (2010).

61. Stegle, O., Parts, L., Piipari, M., Winn, J. & Durbin, R. Using probabilistic estimation of expression residuals (PEER) to obtain increased power and interpretability of gene expression analyses. *Nat. Protoc.* **7**, 500–507 (2012).

62. Ongen, H., Buil, A., Brown, A. A., Dermitzakis, E. T. & Delaneau, O. Fast and efficient QTL mapper for thousands of molecular phenotypes. *Bioinformatics* **32**, 1479–1485 (2016).

63. Wang, G., Sarkar, A., Carbonetto, P. & Stephens, M. A Simple New Approach to Variable Selection in Regression, with Application to Genetic Fine Mapping. *J. R. Stat. Soc. Series B Stat. Methodol.* **82**, 1273–1300 (2020).

64. Ehsan, N. *et al.* Haplotype-aware modeling of cis-regulatory effects highlights the gaps remaining in eQTL data. *bioRxiv* 2022.01.28.478116 (2022) doi:10.1101/2022.01.28.478116.

65. Petrovski, S., Wang, Q., Heinzen, E. L., Allen, A. S. & Goldstein, D. B. Genic intolerance to functional variation and the interpretation of personal genomes. *PLoS Genet.* **9**, e1003709 (2013).

66. Karczewski, K. J. *et al.* The mutational constraint spectrum quantified from variation in 141,456 humans. *Nature* **581**, 434–443 (2020).

67. Collins, R. L. *et al.* A cross-disorder dosage sensitivity map of the human genome. *Cell* **185**, 3041–3055.e25 (2022).

68. Agarwal, I., Fuller, Z. L., Myers, S. R. & Przeworski, M. Relating pathogenic loss-of-function mutations in humans to their evolutionary fitness costs. *Elife* **12**, (2023).

69. ENCODE Project Consortium *et al.* Expanded encyclopaedias of DNA elements in the human and mouse genomes. *Nature* **583**, 699–710 (2020).

70. Lawrence, M. *et al.* Software for computing and annotating genomic ranges. *PLoS Comput. Biol.* **9**, e1003118 (2013).

71. Quinlan, A. R. & Hall, I. M. BEDTools: a flexible suite of utilities for comparing genomic features. *Bioinformatics* **26**, 841–842 (2010).

72. Schmidt, E. M. *et al.* GREGOR: evaluating global enrichment of trait-associated variants in epigenomic features using a systematic, data-driven approach. *Bioinformatics* **31**, 2601–2606 (2015).

73. Ernst, J. & Kellis, M. ChromHMM: automating chromatin-state discovery and characterization. *Nat. Methods* **9**, 215–216 (2012).

74. Biddanda, A., Rice, D. P. & Novembre, J. A variant-centric perspective on geographic patterns of human allele frequency variation. *Elife* **9**, (2020).

75. Wen, X., Lee, Y., Luca, F. & Pique-Regi, R. Efficient Integrative Multi-SNP Association Analysis via Deterministic Approximation of Posteriors. *Am. J. Hum. Genet.* **98**, 1114–1129 (2016).

76. Marcus, J. H. & Novembre, J. Visualizing the geography of genetic variants. *Bioinformatics* **33**, 594–595 (2017).

77. Garrison, E., Kronenberg, Z. N., Dawson, E. T., Pedersen, B. S. & Prins, P. A spectrum of free software tools for processing the VCF variant call format: vcflib, bio-vcf, cyvcf2, hts-nim and slivar. *PLoS Comput. Biol.* **18**, e1009123 (2022).
